# Dedifferentiation unlocks keratinocyte competence for metaplasia and tumorigenesis in the foregut

**DOI:** 10.64898/2026.06.24.734259

**Authors:** Alizée Vercauteren Drubbel, Sheleya Pirard, Benjamin Beck

**Author notes:** Co-corresponding authors &.

## Abstract

How tissue injury shapes cell competence to undergo malignant transformation remains poorly understood. Esophageal metaplasia, a precancerous lesion driven by chronic acid reflux, can arise from the conversion of squamous progenitors into a columnar-like state, but the factors governing this plasticity remain unclear. Here we show that GATA4, overexpressed in esophageal metaplasia and adenocarcinoma, drives columnar metaplasia in squamous progenitors at the squamo-columnar junction but is insufficient, and even toxic, in keratinocytes outside this region. Using inducible transgenic mouse models, we find that reactivation of Hedgehog signaling expands the pool of progenitors permissive to GATA4-mediated reprogramming, driving gastric-like metaplasia even within the esophagus. Combined Hedgehog activation and GATA4 expression further induce adenosquamous-like neoplasms and stromal and immune remodeling reminiscent of the metaplastic microenvironment. Since Hedgehog signaling is reactivated by gastroesophageal reflux, chronic injury may generate a field of dedifferentiated progenitors poised for malignant progression upon oncogene acquisition. These findings demonstrate that a prior cell state transition, induced by environmental injury, can unlock oncogenic competence, establishing a mechanistic framework linking epithelial plasticity, developmental transcription factor reactivation, and lineage-specific cancer susceptibility with broad implications for precancerous metaplastic states.

## Introduction

Tumorigenesis is a multistep process driven by both genetic and phenotypic changes within epithelial tissues. Classical carcinogenesis models posited that oncogenic mutations accumulate in individual cells and subsequently expand under the influence of extrinsic stimuli such as inflammation, tissue injury, or environmental insults^1^. A major advance in our understanding of early tumor evolution has come from sequencing studies showing that normal epithelial tissues frequently harbor numerous oncogenic mutations. In the human epidermis and esophagus, nearly all individuals carry thousands of clones containing driver mutations such as *TP53* or *NOTCH1*^2,3^. Yet, only a minority of these clones progress to cancer, underscoring that mutational burden alone is insufficient for malignant transformation. Instead, the emergence of neoplasia appears to depend on the cellular context in which mutations arise, as well as the state of the surrounding microenvironment^4^.

The esophagus provides an instructive model for dissecting these interactions. Indeed, the healthy esophageal epithelium contains abundant driver-mutant clones^3^, and two major cancer types commonly arise: esophageal squamous cell carcinoma (eSCC) and esophageal adenocarcinoma (eAC). The incidence of eAC has risen dramatically in Western countries and is strongly linked to chronic gastroesophageal reflux disease (GERD), obesity, and esophagitis^5^. These risk factors promote the development of Barrett’s esophagus (BE), a metaplastic conversion of the stratified squamous epithelium into a columnar epithelium resembling gastric or intestinal mucosa^6^. Metaplasia is a conserved injury response in multiple organs, including the pancreas, lung, and stomach, in which one differentiated cell type is replaced by another^7^. In the esophagus, BE arises near the squamo-columnar junction (SCJ), where gastric and esophageal epithelia meet. Gastric progenitors from the SCJ can be at the origin of columnar metaplasia^8^, and in most human cases, BE frequently shares somatic mutations with adjacent gastric mucosa^9^. However, studies also indicate that esophageal squamous keratinocytes, particularly those at the SCJ, retain a degree of plasticity. Forced expression of CDX2^10^, loss of lineage-specifying transcription factor^11^ or BMP pathway activation^12^ can induce aspects of intestinal differentiation in murine squamous progenitors, though only in discrete subpopulations, suggesting that metaplasia may have heterogeneous origins^13^. In line with this hypothesis, a recent study showed that BE are polyclonal lesions where lineages connecting BE to both esophageal and gastric tissues can be identified^14^. But, to date, no *in vivo* evidence has shown that squamous cells located distant from the SCJ retain the capacity to undergo true metaplastic reprogramming.

Our previous work revealed that chronic acid reflux reactivates Hedgehog (HH) signaling in esophageal progenitors, inducing dedifferentiation into a transcriptionally and epigenetically embryonic-like state^15^. This state resembles early foregut progenitors, which are bipotent during development and capable of generating either squamous or columnar epithelium^16^. Importantly, these dedifferentiated cells can persist for extended periods, raising the possibility that they constitute a reservoir poised for metaplastic transformation^17^.

Developmental transcription factors that regulate early foregut patterning^18^ are likely critical mediators of such lineage reprogramming. Among these, GATA4 is essential for gastric and hindgut development and represses squamous identity while promoting columnar differentiation^19–21^ (**Extended Data Fig. 1a**). Moreover, GATA4 is aberrantly expressed in BE^21^ and is the most frequently amplified transcription factor in eAC^22^. By contrast, *GATA4* amplification is rare in eSCC, suggesting that its activation confers selective advantage in columnar, but not squamous, lineages. Yet, the effects of GATA4 overexpression in uninjured esophageal squamous epithelium remain unknown.

In this study, we tested the hypothesis that GATA4 expression is sufficient to induce columnar metaplasia when introduced into foregut progenitors. Using inducible transgenic mouse models, we found that long-term GATA4 expression had minimal effect on gastric mucosa but induced columnar-like metaplasia at the SCJ when expressed in squamous progenitors. In contrast, GATA4-positive esophageal keratinocytes were rapidly eliminated unless they had first undergone HH-induced dedifferentiation. When introduced into these HH-reactivated cells, GATA4 drove the formation of metaplastic clones with gastric features at distance from the SCJ including within the esophagus, and even neoplasms of mixed-histology.

Together, these findings demonstrate that cell state transitions, not oncogene activation alone, determine the competence of esophageal progenitors to undergo metaplasia and malignant progression. This work provides a mechanistic framework linking chronic injury, epithelial plasticity, and lineage-specific oncogenic susceptibility in the foregut.

## Results

### GATA4 expression in squamous lineage induces columnar metaplasia

In mouse, gastric transcription factor GATA4 is virtually absent from the esophageal squamous epithelium, as demonstrated by immunostaining (**Fig. 1a and Extended Data Fig. 1a, b**). Analysis of cancer genomics datasets revealed that *GATA4* is frequently amplified in esophageal adenocarcinoma (eAC), and that these amplifications are associated with increased *GATA4* mRNA expression (**Fig. 1b, c and Extended Data Fig. 1c**). Immunostaining and analysis of transcriptomic data further showed that *GATA4* expression is already elevated in human Barrett’s esophagus (BE), the premalignant precursor lesion of eAC (**Fig. 1d, e and Extended Data Fig. 1d**), consistent with previously published data^21^.

**Figure 1.**
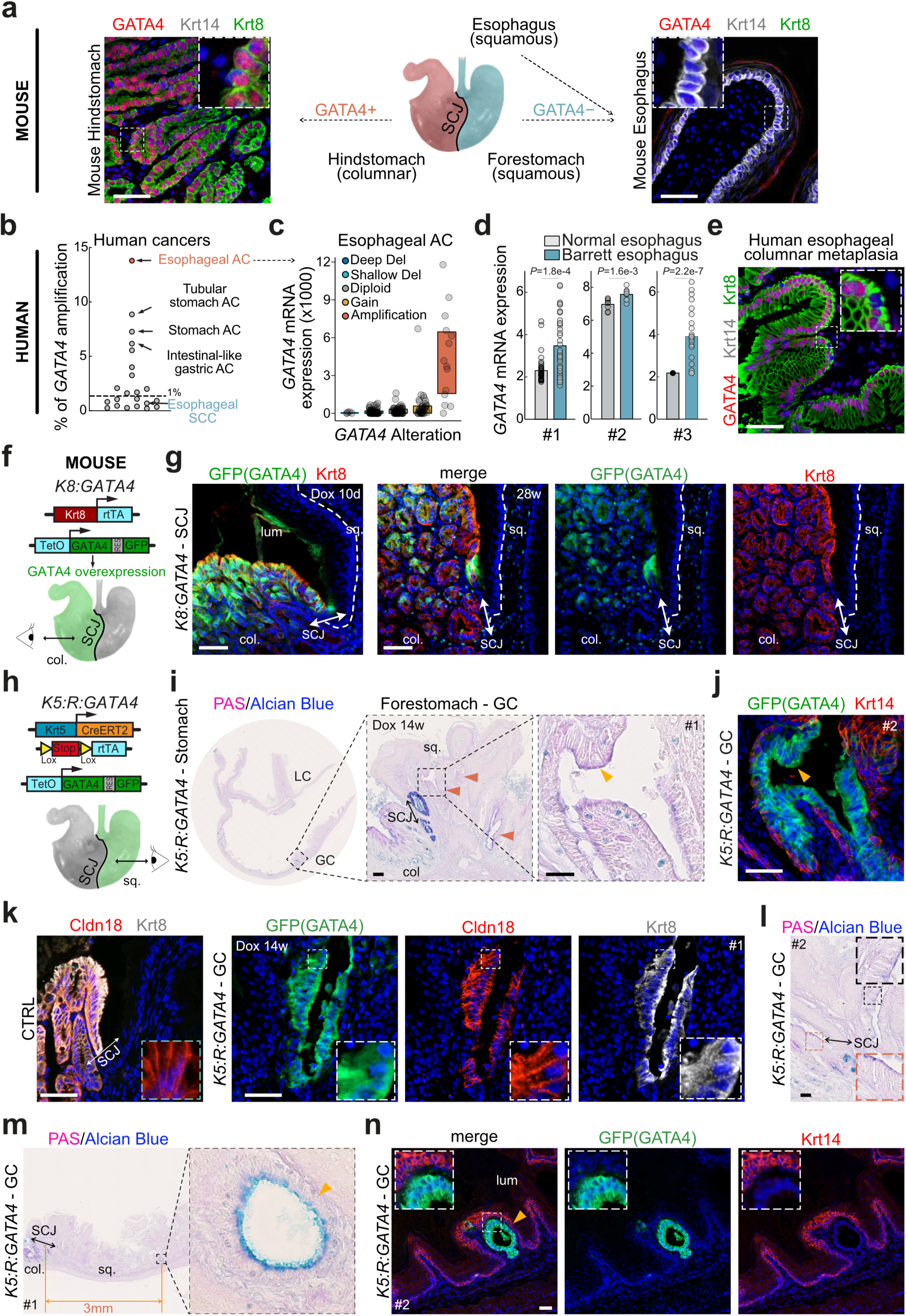
GATA4 overexpression drives columnar metaplasia development from squamous progenitors at the squamo-columnar junction. **a.** Immunostaining for GATA4, Krt14, and Krt8 in mouse stomach corpus and esophagus. **b.** Dot plot showing the incidence of *GATA4* gene amplification across human cancers. Esophageal adenocarcinoma (AC), gastric AC, and esophageal squamous cell carcinoma (SCC) are highlighted. **c.** Box plot showing the distribution of *GATA4* copy number alterations in human esophageal AC. **d.** Histogram showing *GATA4* mRNA expression in normal esophagus and Barrett’s esophagus samples. Data are from three independent studies (GSE39491, *n*=161; GSE34619, *n*=36; GSE36223, *n*=93 per condition). **e.** Immunostaining for GATA4, Krt14, and Krt8 in a representative human Barrett’s esophagus sample. **f.** Genetic model used to overexpress GATA4 in the gastric columnar epithelium (*Krt8-rtTA:TetO-GATA4-IRES-GFP*, hereafter referred to as K8:GATA4). **g.** Immunostaining for GFP (transgenic GATA4) and Krt8 in mouse stomach corpus after 10 days or 28 weeks of doxycycline (Dox) treatment. **h.** Genetic model used to overexpress GATA4 in the esophageal and gastric squamous epithelium (forestomach) (*Krt5-CreER:Rosa26-rtTA:TetO-GATA4-IRES-GFP*, hereafter referred to as K5:R:GATA4). **i.** Periodic Acid–Schiff (PAS)/Alcian blue staining in K5:R:GATA4 mice after 14 weeks of doxycycline treatment. Orange arrows highlight PAS/Alcian blue-positive lesions. **j.** Immunostaining for GFP (GATA4) and Krt14 in mouse forestomach after 14 weeks of doxycycline treatment. Panels (**i**) and (**j**) were performed on serial sections. **k.** Immunostaining for gastric markers Cldn18 and Krt8 at the gastric squamo-columnar junction (SCJ) of control (wild-type, CTRL) mice and the greater curvature of forestomach of K5:R:GATA4 mice after 14 weeks of doxycycline treatment, together with GFP (transgenic GATA4) staining. **l.** Periodic Acid–Schiff (PAS)/Alcian blue staining in K5:R:GATA4 mice after 14 weeks of doxycycline treatment. Panels (**k**) and (**l**) were performed on serial sections. **m.** Periodic Acid–Schiff (PAS)/Alcian blue staining in K5:R:GATA4 mice after 14 weeks of doxycycline treatment, showing metaplastic lesions distant from the SCJ at the greater curvature of forestomach. **n.** Immunostaining for GFP (GATA4) and Krt14 in mouse forestomach after 14 weeks of doxycycline treatment. Panels (**m**) and (**n**) were performed on serial sections. In all panels, nuclear Hoechst staining is shown in blue. Scale bars = 50 µm unless otherwise stated. Sq, squamous; col, columnar; SCJ, squamo-columnar junction; lum, lumen; LC, lesser curvature; GC, greater curvature.

To investigate the functional consequences of GATA4 overexpression in epithelial lineages, we generated a doxycycline-inducible transgenic mouse model co-expressing GATA4 and a GFP reporter (TetO-GATA4-GFP). These mice were crossed with a Krt8 driver line to target columnar epithelial lineage (**Extended Data Fig. 1e-g**). *Krt8-rtTA:TetO-GATA4-GFP* mice (hereafter K8:GATA4) were treated with doxycycline for up to 28 weeks to trigger GATA4 overexpression (**Fig. 1f**). As expected, GFP-positive cells were detected in the gastric mucosa, but not in the squamous epithelium (**Fig. 1g and Extended Data Fig. 1g**). Despite sustained GATA4 overexpression for more than six months, no major histological or phenotypic alterations were observed in the stomach (**Fig. 1g and Extended Data Fig. 2a-b**).

To investigate the functional consequences of GATA4 overexpression in squamous epithelial lineage, we crossed TetO-GATA4-GFP mice with a Krt5 driver line. To bypass potential transgene silencing associated with loss of Krt5 expression, we generated *Krt5-CreER:Rosa26-rtTA:TetO-GATA4-GFP* mice (hereafter K5:R:GATA4), in which sustained GATA4 expression is maintained independently of squamous cell identity (**Fig. 1h and Extended Data Fig. 2c, d**). After 14 weeks of GATA4 induction in the squamous lineage of K5:R:GATA4 mice, we observed histological modifications at the greater curvature (GC) of the forestomach (Extended Data Fig. 2e). These included the appearance of periodic acid-Schiff (PAS)-positive structures in the vicinity of the squamo-columnar junction (SCJ) (**Fig. 1i**). Staining of serial sections revealed that these structures were GFP-positive, confirming their squamous origin (**Fig. 1j and Extended Data Fig. 3a**). Furthermore, a subset of GFP-positive clones expressed the gastric markers Krt8 and Claudin18 but not the intestinal markers Cdx2 (**Fig. 1k and Extended Data Fig. 3b-d**) and displayed a morphology reminiscent of SCJ columnar cells (**Fig. 1l**), consistent with transdifferentiation of a subset of foregut squamous progenitors. Notably, some PAS/Alcian blue-positive, GATA4-GFP positive, Krt14-negative structures were entirely embedded within the squamous epithelium and located several millimeters away from the SCJ, indicating that rare squamous progenitors outside the immediate SCJ region can undergo transdifferentiation upon GATA4 overexpression (**Fig. 1m, n and Extended Data Fig. 3e**).

### Hedgehog pathway activation promotes reprogramming upon GATA4 overexpression

Keratinocytes of the SCJ are characterized by constitutive activation of the Hedgehog (HH) signaling pathway and exhibit an embryonic-like phenotype^15,23^. Notably, HH pathway activation alone is sufficient to induce a transcriptomic and epigenetic embryonic program in esophageal progenitor cells *in vivo*^15^. These observations led us to hypothesize that HH pathway activation may facilitate GATA4-induced cell reprogramming. To test this, we generated the *Krt5-CreER:Rosa26-SmoM2:Rosa26-rtTA:TetO-GATA4-GFP* (K5:Smo:R:GATA4) model, which enables tamoxifen-inducible lineage tracing of SmoM2-positive esophageal cells harboring constitutive HH activation, combined with doxycycline-inducible GATA4 overexpression (**Fig. 2a**).

**Figure 2.**
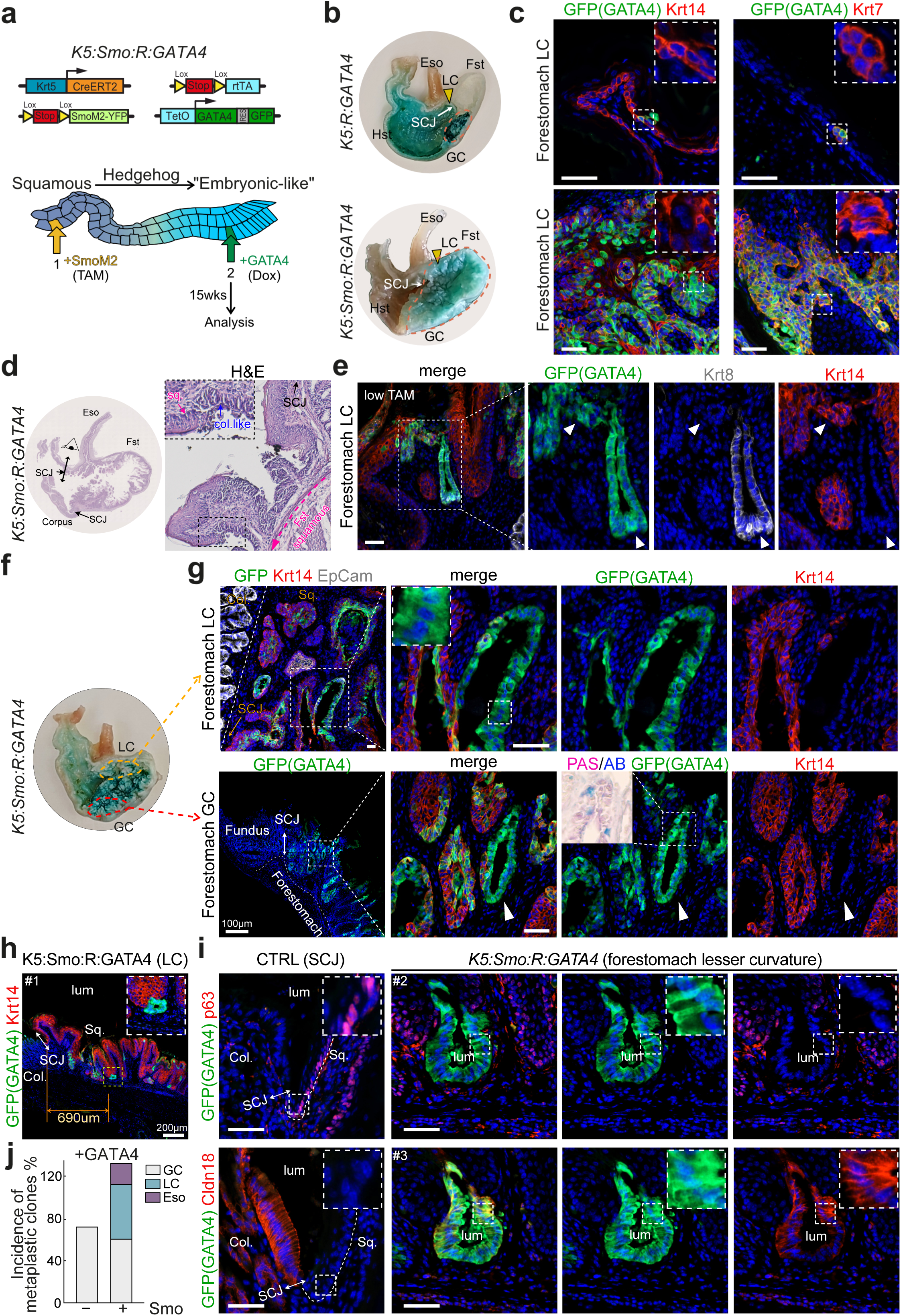
Hedgehog pathway activation promotes GATA4-driven columnar metaplasia development from squamous progenitors at distance from the squamo-columnar junction. **a.** Experimental design to express GATA4 in Hedgehog-dedifferentiated cells induced by SmoM2 activation using the Krt5-CreER:Rosa26-rtTA:Rosa26-SmoM2:TetO-GATA4-IRES-GFP mouse model, hereafter referred to as K5:Smo:R:GATA4. All K5:Smo:R:GATA4 mice shown in this figure were fed doxycycline (Dox) for 14 weeks (GATA4 expression), starting 1 week after tamoxifen (TAM) induction (Hedgehog activation). **b.** Macroscopic image of the stomach of K5:R:GATA4 and K5:Smo:R:GATA4 mice fed a doxycycline diet containing blue dye for 14 weeks following 1 week of TAM induction. Of note, the blue dye colors certain regions of the stomach independently of GATA4 expression. **c.** Immunostaining for GFP (GATA4) and Krt14 or Krt7 in the lesser curvature of forestomach of K5:R:GATA4 and K5:Smo:R:GATA4 mice 1 week after TAM followed by 14 weeks Dox induction. **d.** Hematoxylin / Eosin (H/E) staining in K5:Smo:R:GATA4 mouse forestomach showing columnar-like cells distant from the SCJ. **e.** Immunostaining for GFP (GATA4), Krt8 and Krt14 in the greater curvature of forestomach of K5:Smo:R:GATA4 mice after low-dose TAM (1mg) induction followed by doxycycline treatment. White arrows indicate a squamous region and a columnar metaplastic region, both arising from a common GFP+ (GATA4) Krt5+ squamous progenitor. **f.** Macroscopic image of the stomach of K5:Smo:R:GATA4 mice fed a doxycycline diet containing blue dye following TAM induction. **g.** Immunostaining for GFP (GATA4) and Krt14 in K5:Smo:R:GATA4 mouse forestomach lesser curvature and greater curvature. Inset shows Periodic Acid–Schiff (PAS)/Alcian blue (AB) staining of the indicated area. **h.** Immunostaining for GFP (GATA4) and Krt14 in K5:Smo:R:GATA4 mouse forestomach lesser curvature, showing metaplastic lesions at distance from the SCJ. **i.** Immunostaining for GFP (GATA4) and p63 or Cldn18 in K5:Smo:R:GATA4 mouse forestomach lesser curvature. Left panels show the expression pattern at the SCJ in control animals. Panels (**h**) and (**i**) were performed on serial sections. **j.** Bar plot showing the distribution of metaplastic lesions in the greater curvature (GC), lesser curvature (LC), and distal esophagus (Eso) in K5:R:GATA4 and K5:Smo:R:GATA4 animals. In all panels, nuclear Hoechst staining is shown in blue. Scale bars = 50 µm unless otherwise stated. Sq, squamous; col, columnar; SCJ, squamo-columnar junction; lum, lumen; LC, lesser curvature; GC, greater curvature.

Strikingly, whereas 14 weeks of doxycycline treatment in K5:R:GATA4 animals resulted in the appearance of PAS-positive metaplastic clones in the SCJ area of the GC and only few GATA4+ scattered cells in the lesser curvature (LC) and the esophagus, the same treatment in K5:Smo:R:GATA4 mice led to the development of neoplastic lesions in the whole forestomach (**Fig.2b, c and Extended Data Fig. 4a, b**). These lesions were characterized by heterogeneous clusters of GFP/Krt7 double-positive cells expressing variable levels of the squamous marker Krt14 (**Fig. 2c and Extended Data Fig. 4b,c**). Hematoxylin and eosin (H&E) staining revealed the coexistence of squamous-like and columnar-like cell populations intermingled within the same region (**Fig. 2 d and Extended Data Fig. 4c**). Similar mixed structures were also observed following recombination with low doses of tamoxifen, revealing that individual progenitor cells possess bipotent differentiation potential (**Fig. 2e**). The coexistence of squamous and columnar-like cells in the foregut is reminiscent of adenosquamous carcinoma, a rare histological subtype of cancer associated with chronic gastroesophageal reflux, whose cellular origin remains poorly understood^24,25^.

Interstingly, in K5:Smo:R:GATA4 animals greater and lesser curvatures both showed similar GFP columnar-like structures, some positive for Krt14 and other negative for Krt14 and positive for Alcian blue (**Fig. 2f, g**). Presence of transdifferentiated cells at the LC was likely the consequence of the maintenance of GATA4 overexpressing cells in SmoM2+ embryonic-like progenitors (**Extended Data Fig. 4d**).

GFP-positive, Krt14-negative structures were also identified far from the SCJ at the lesser curvature, a region that remained devoid of metaplastic lesions in K5:R:GATA4 animals (**Fig. 2h-i**). Further characterization of these structures revealed loss of the basal squamous markers p63 and Krt14, alongside induction of gastric markers including Cldn18, Krt8 and Krt7, supporting a complete transdifferentiation process and the formation of metaplasia-like lesions (**Fig. 2i and Extended Data Fig. 5a-d**). Notably, these clones did not express Cdx2, indicating transdifferentiation toward a gastric-like rather than intestinal-like epithelial identity (**Extended Data Fig. 5e**). Of note, similar clones were also identified in the distal esophagus of K5:Smo:R:GATA4 mice (**Fig. 2j**), an organ previously reported to be resistant to transdifferentiation^10,26^. Collectively, these results demonstrate that HH pathway activation facilitates squamous progenitor transdifferentiation, and that sustained GATA4 overexpression in this context is sufficient to drive gastric-like metaplasia as well as adenosquamous-like tumor formation.

### Epithelial HH/GATA4 activation drives coordinated stromal and immune reprogramming

To further characterize the consequences of GATA4 expression in HH-positive squamous progenitors, we dissected the lesser and greater curvatures of the forestomach from control and K5:Smo:R:GATA4 animals and performed single-cell RNA sequencing (scRNA-seq) across conditions. This analysis confirmed the presence of *Krt14*/*Krt7* double-positive cells at both the greater and lesser curvatures exclusively in the K5:Smo:R:GATA4 condition (**Fig. 3a and Extended Data Fig.6a-c**). Similar transitional (squamous/columnar) basal progenitors (*Krt14*+/*Krt7*+) were also observed in human Barrett’s esophagus analysed with single-cell transcriptomics^9,27^.

**Figure 3.**
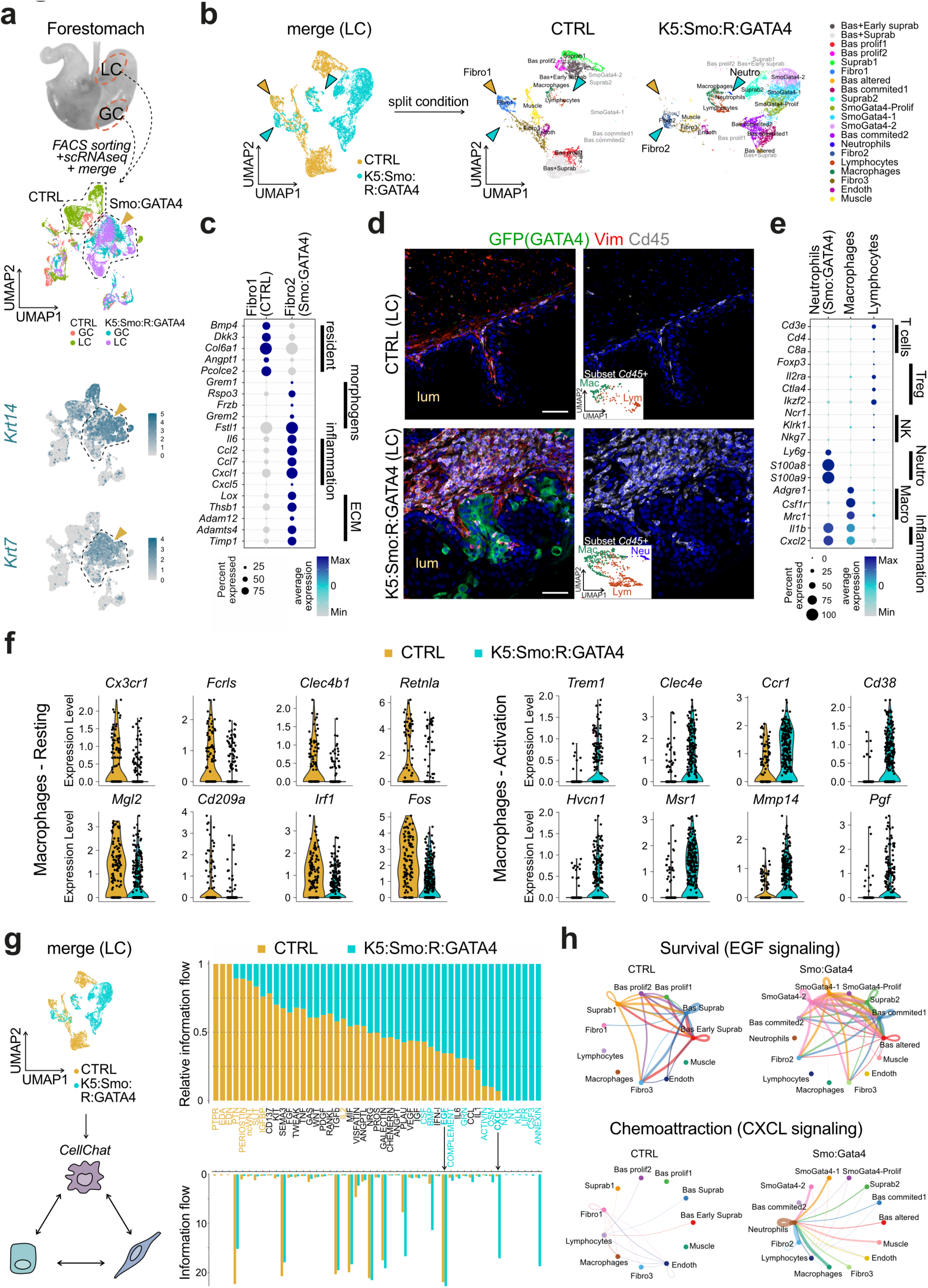
Hedgehog pathway activation promotes GATA4-driven gastric reprogramming from esophageal progenitors. **a.** UMAP from merged single-cell RNA sequencing (scRNA-seq) data of cells from the greater and lesser curvature of K5:Smo:R:GATA4 and control (CTRL) mice. Cells are colored by condition. Feature plots depicting *Krt14* and *Krt7* mRNA expression highlight the presence of double-positive cells in both regions of K5:Smo:R:GATA4 (yellow arrows). **b.** Annotated UMAP of merged scRNA-seq data from lesser curvature cells of the forestomach of CTRL and K5:Smo:R:GATA4 mice, showing changes in stromal cell composition (yellow and blue arrows). **c.** Dot plot representing expression of fibroblast markers in the two fibroblast cell clusters (Fibro1- CTRL and Fibro2 - Smo:GATA4) from UMAP depicted in (**b**). **d.** Immunostaining for GFP (GATA4), Vimentin and CD45 in CTRL and K5:Smo:R:GATA4 mouse forestomach lesser curvature. Nuclear Hoechst staining is shown in blue; lum, lumen. Scale bars = 50 µm. Insets show the UMAP representation of immune cell clusters from (**b**). **e.** Dot plot representing expression of immune markers in the three immune cell clusters (Neutrophils- Smo:GATA4, Marcophages and Lymphocytes) from UMAP depicted in (**b**). **f.** Violin plot representing the expression of resting- or activated- macrophages in the macrophage cell cluster of CTRL and Smo:GATA4 conditions, extracted from the dataset shown in (**b**). **g.** Relative information flow measured with *CellChat* in CTRL and K5:Smo:R:GATA4 mouse lesser curvature forestomach cells depicted in (**b**), showing alterations in intercellular signaling when SmoM2 and GATA4 are co-expressed in epithelial cells. Non-significant pathways are shown in black. **h.** Circle plots illustrating modifications of EGF, and CXCL signaling pathway networks in K5:Smo:R:GATA4 mouse forestomach compared to CTRL. All K5:Smo:R:GATA4 mice shown in this figure were fed doxycycline (Dox) for 14 weeks (GATA4 expression), starting 1 week after tamoxifen (TAM) induction (Hedgehog activation). Uniform Manifold Approximation and Projection = UMAP.

We then focused our scRNA-seq analysis on the lesser curvatures (**Fig. 3b** and **Extended Data Fig. 7a-c)** and confirmed that some epithelial cell clusters co-expressing YFP (*SmoM2*) and GFP (*GATA4*) displayed several markers of gastric lineage (*Cldn18*, *Gkn3*, *Tff1*, *Ctse*) but not of intestinal lineage (*Gata6*, *Cdx2*, *Muc2*) (**Extended Data Fig. 7d–f**), confirming that HH/GATA4-driven reprogramming directs progenitors toward a gastric rather than intestinal columnar identity. Analysis of the lesser curvature revealed extensive microenvironmental remodeling upon HH/GATA4 expression, including the emergence of a pro-inflammatory fibroblast population (cluster Fibro2) expressing *Il6*, *Ccl2*, *Ccl7*, and *Cxcl1*, as well as extracellular matrix remodeling components (*Lox*, *Thbs1*, *Adam12*, *Adam14*, and *Timp1*), and morphogenes (*Grem1*, *Grem2* and *Rspo3*) (**Fig. 3c**).

Immune cell profiling by scRNA-seq showed an increased relative proportion of macrophages, lymphocytes, and neutrophils, consistent with a higher density of CD45+ cells observed by immunofluorescence in forestomach from K5:Smo:R:GATA4 animals (**Fig. 3d, e and Extended Data Fig. 7c**). Neutrophils (*Ly6g*+/*Csf3r*+/*S100a8*+/*S100a9*+; cluster 19) were detected almost exclusively in the transgenic condition, suggesting active inflammatory recruitment (**Fig. 3d and Extended Data Fig. 7c**). In contrast, the lymphocyte compartment (*Cd3e*+ ; cluster 13) displays a regulatory T-cell–like signature, with prominent expression of *Foxp3*, *Il2ra*, *Ctla4*, and *Ikzf2*, suggesting expansion of an immunoregulatory population rather than a cytotoxic/NK-like response (**Fig. 3e**). Together, these data indicate that immune remodeling in the transgenic tissue is primarily driven by inflammatory myeloid recruitment and activation, accompanied by a likely compensatory expansion of regulatory T cells.

In the K5:Smo:R:GATA4 condition, cluster 14 (*Adgre1*+/*Csf1r*+/*Mrc1*+ ; macrophages) exhibited a reduction in resident/homeostatic markers (*Cx3cr1*, *Retnla*, *Fos*) and an enrichment of genes associated with activation, tissue remodeling, and immunoregulation (*Trem1*, *Clec4e*, *Ccr1*, *Cd38*, *Hvcn1*, *Msr1*, *Mmp14*, *Pgf*), consistent with an activated macrophage state with remodeling and immunoregulatory features (**Fig. 3f**).

These findings support a model in which the transgenic condition promotes inflammatory myeloid recruitment and macrophage reprogramming, thereby establishing a tissue-remodeling immune niche consistent with chronic injury, aberrant repair, and metaplastic progression, as observed in Barrett’s esophagus^27^. These changes are further associated with increased EGF and CXCL signaling (**Fig. 3g, h**), consistent with stromal reprogramming described in nascent esophageal tumors^28^ and human Barrett’s metaplasia^27^ respectively.

### GATA4 expression promotes a gastric lineage program in a subset of esophageal squamous progenitors

Since esophageal progenitors had been previously described as resistant to transdifferentiation *in vivo*, we then further investigated the fate of GATA4-positive esophageal progenitors.

First, we observed that following HH pathway activation, GATA4-expressing cells were also maintained as clusters of Krt8-positive epithelial cells within the squamous esophageal mucosa, while being delaminated from their SmoM2-negative counterparts (**Fig. 4a, b**).

**Figure 4.**
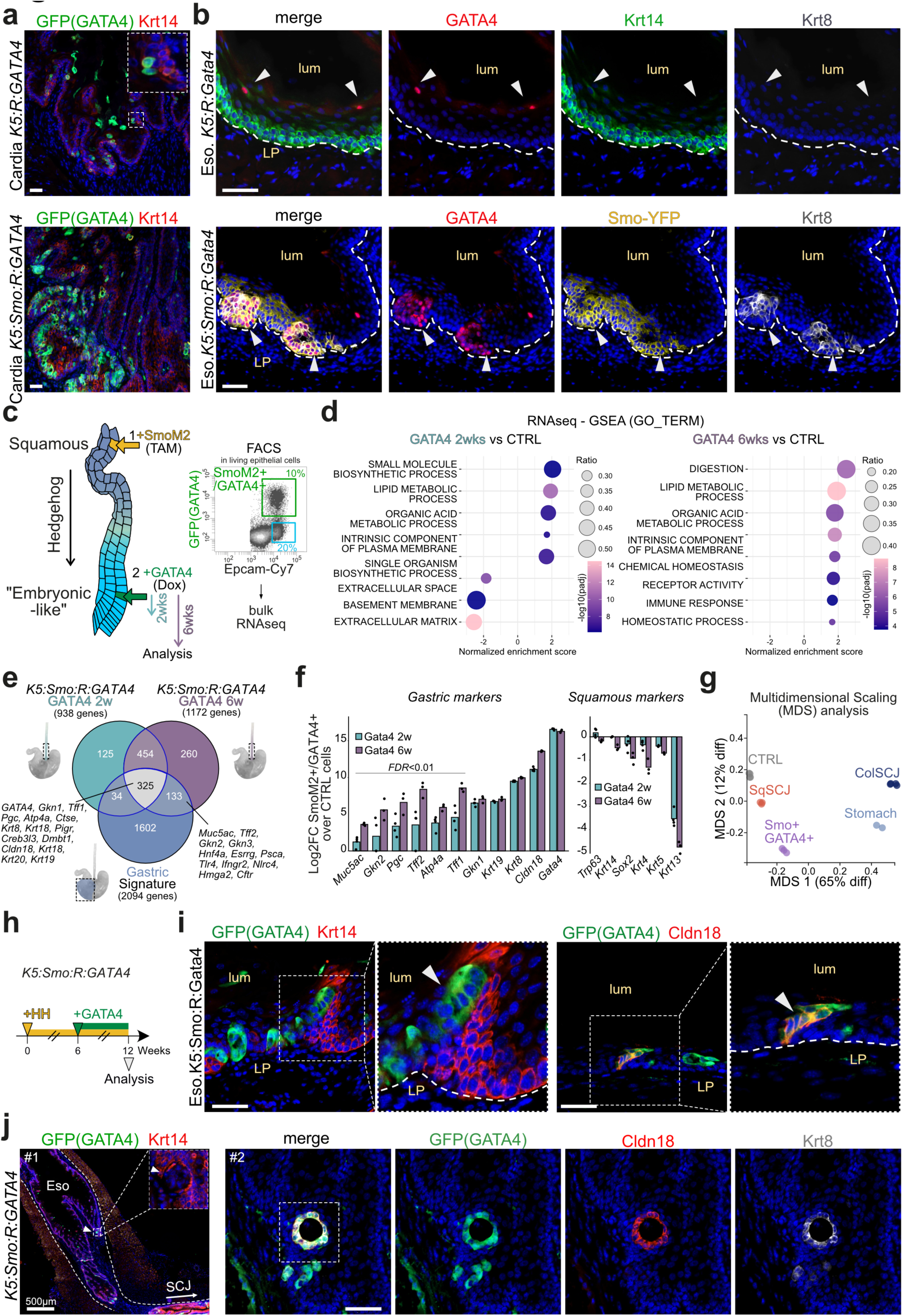
Hedgehog pathway activation promotes GATA4-driven gastric reprogramming from esophageal progenitors. **a.** Immunostaining for GFP (GATA4) and Krt14 in the esophageal cardia region of K5:R:GATA4 and K5:Smo:R:GATA4 mice after Tamoxifen (TAM) induction followed by doxycycline (Dox) treatment. **b.** Immunostaining for GATA4, Krt14, and Krt8 in K5:R:GATA4 mouse esophagus (top), or for GATA4, SmoM2, and Krt8 in K5:Smo:R:GATA4 mouse esophagus (bottom). Samples were analyzed following TAM and Dox induction. **c.** Experimental design to induce GATA4 expression for 2 or 6 weeks starting 6 weeks after HH pathway activation (SmoM2) in esophageal cells. FACS strategy to isolate SmoM2+ (Epcam High) GFP-labeled GATA4-overexpressing cells (SmoM2+/GATA4+). **d.** Dot plot summarizing GSEA results from GATA4+ cells compared to control esophageal cells after 2 or 6 weeks of Dox treatment, design depicted in (**c**). **e.** Venn diagram showing the overlap between genes upregulated in GATA4+ cells after 2 or 6 weeks of Dox treatment and genes upregulated in gastric columnar epithelial cells compared to control esophageal squamous progenitors. **f.** Bar plot summarizing the expression of selected gastric and squamous markers in GATA4+ cells after 2 or 6 weeks of doxycycline treatment. *FDR* values were calculated using the Benjamini–Hochberg method in DESeq2 when comparing 2- and 6-week doxycycline treatments. **g.** Multidimensional scaling (MDS) plot showing the distribution of RNA profiles from control, junction keratinocytes, SmoM2+/GATA4+, gastric columnar, and junction columnar cells, as measured by bulk RNA sequencing. **h.** Experimental design to study the impact of GATA4 overexpression in HH-dedifferentiated esophageal progenitors *in vivo*. **i.** Immunostaining for GFP (GATA4) and Krt14 or Cldn18 in K5:Smo:R:GATA4 mouse esophagus after 6 weeks of Dox treatment starting 6 weeks after TAM induction. **j.** Immunostaining for GFP (GATA4), Krt14 or GFP (GATA4), Cldn18 and Krt8 in K5:Smo:R:GATA4 mouse esophagus after 14 weeks of Dox treatment. Krt14 and Cldn18 stainings were performed on serial sections. In all panels, nuclear Hoechst staining is shown in blue. Scale bars = 50 µm unless otherwise stated. LP, lamina propria; lum, lumen; Eso, esophagus; SCJ, squamo-columnar junction; SqSCJ, Squamous cells from de SCJ; ColSCJ, Columnar cells from de SCJ.

To determine how esophageal cells could be reprogrammed *in vivo*, we examined different time points in K5:Smo:R:GATA4 animals. As previously reported, squamous marker expression progressively decreases following SmoM2 activation^15^. We therefore focused our first analyses on a protocol in which cells were first dedifferentiated for six weeks prior to GATA4 induction, a stage at which they already exhibited embryonic-like features.

We performed transcriptomic profiling of FACS-sorted SmoM2+/GATA4+ cells from K5:Smo:R:GATA4 mice by bulk RNA sequencing at two and six weeks after GATA4 induction (**Fig. 4c**). Gene set enrichment analysis (GSEA) on the genes deregulated in SmoM2+/GATA4+ compared to CTRL esophageal cells revealed enrichment for biological processes related to “*digestion*” and “*lipase activity*” (**Fig. 4d and Extended Data Fig. 8a**). GSEA of genes deregulated 6 weeks relative to 2 weeks after GATA4 induction, revealed enrichment for gene sets involved in “extracellular matrix” and “*immune response*” (**Extended Data Fig. 8c**). Comparison of upregulated genes against reference transcriptomic signatures from distinct digestive tract lineages^15^ showed that GATA4-

GFP-positive cells were enriched for gastric epithelial markers such as *Gkn1*, *Pgc*, *Atp4a* and *Cldn18* (**Fig. 4e and Extended Data Fig. 8d**). Importantly, this gastric program was induced rapidly : 38% and 39% of upregulated transcripts at two and six weeks of treatment, respectively, overlapped with gastric epithelial signatures (**Fig. 4e and Extended Data Fig. 8b**), indicating that GATA4 promptly initiates a progressive gastric transcripomic program. Analysis of selected genes confirmed the strong upregulation of canonical gastric markers, including *Cldn18*, *Gkn1*, *Atp4a*, *Gkn2*, and *Muc5ac*, alongside the downregulation of squamous lineage markers such as *Krt13*, *Krt5*, *Krt4*, *Krt14*, and *Sox2* (**Fig. 4f**). Several gastric markers including *Tff2*, *Pgc*, *Muc5ac*, and *Gkn2* were expressed at significantly higher levels at six weeks than at two weeks of GATA4 overexpression (**Fig. 4f**), indicating that reprogramming deepens with prolonged GATA4 activity. Multidimensional Scaling (MDS) confirmed that these reprogrammed cells are transcriptionally more similar to gastric cells than to columnar SCJ cells, yet remain distinct from native gastric cells. (**Fig. 4g**).

Of note, a two-week period of HH-induced dedifferentiation was sufficient to permit both the maintenance of GATA4-positive keratinocytes in the esophagus and the upregulation of more than 650 genes upregulated at later time point. GSEA confirmed an enrichment for *digestion* pathway, consistent with the induction of key gastric markers (**Extended Data Fig. 8a-f**). Nonetheless, immunostaining revealed that GATA4 expression in cells dedifferentiated for only two weeks leads to milder phenotype than if cells are dedifferentiated for 6 weeks (**Extended Data Fig. 9a-e**). These results suggest that upon short-term GATA4 expression, the extent of HH-induced dedifferentiation influences the efficiency of morphological remodeling but has a limited impact on the transcriptional program induced by GATA4.

Histological examination of K5:Smo:R:GATA4 esophagi following six weeks of doxycycline treatment revealed phenotypic diversity among GATA4-positive clones (**Fig. 4h**). Some clones retained expression of the squamous marker Krt14, whereas others had lost Krt14 expression and exhibited altered keratinocyte morphology (**Fig. 4i and Extended Data Fig. 9c, d**). Notably, a subset of clones maintained a keratinocyte-like appearance while expressing columnar markers such as Claudin18 (**Fig. 4i and Extended Data Fig. 9e**). At this time point genuine metaplastic clones were not observed in the esophagus, whereas rare metaplastic clones were already detected in the LC of the forestomach, suggesting that esophageal cells are more resistant to metaplasic conversion than forestomach cells (**Fig. 4h, i and Extended Data Fig. 9c-f**). Nevertheless, at later stages, when metaplastic clones were frequently detected at the greater and lesser curvature of the forestomach, we also observed analogue GATA4+/Cldn18+/Krt8+/p63-negative columnar structures in the lower part of the esophagus (**Fig. 4j, 2j and Extended Data Fig. 9g**). These results unambiguously demonstrated that at least some esophageal progenitors retain sufficient plasticity to transdifferentiate *in vivo*. These observations prompted us to further investigate the fate and heterogeneity of GATA4-positive cells at single-cell resolution.

### GATA4 enables dedifferentiated cells to connect squamous and gastric-like lineages

To resolve the cellular dynamics underlying transdifferentiation, we isolated all viable cells from the esophagi of K5:Smo:R:GATA4 mice at the time point when Cldn18+ gastric-like cells first appear, and performed scRNA-seq (**Fig. 5a**). As expected, some cell clusters from the subset of epithelial cells expressed high levels of *Krt8* and *Cldn18* together with lower levels of *Trp63*, the master regulator of squamous differentiation^23^, suggesting they had undergone a full squamous-to-columnar conversion (**Fig. 5b**).

**Figure 5.**
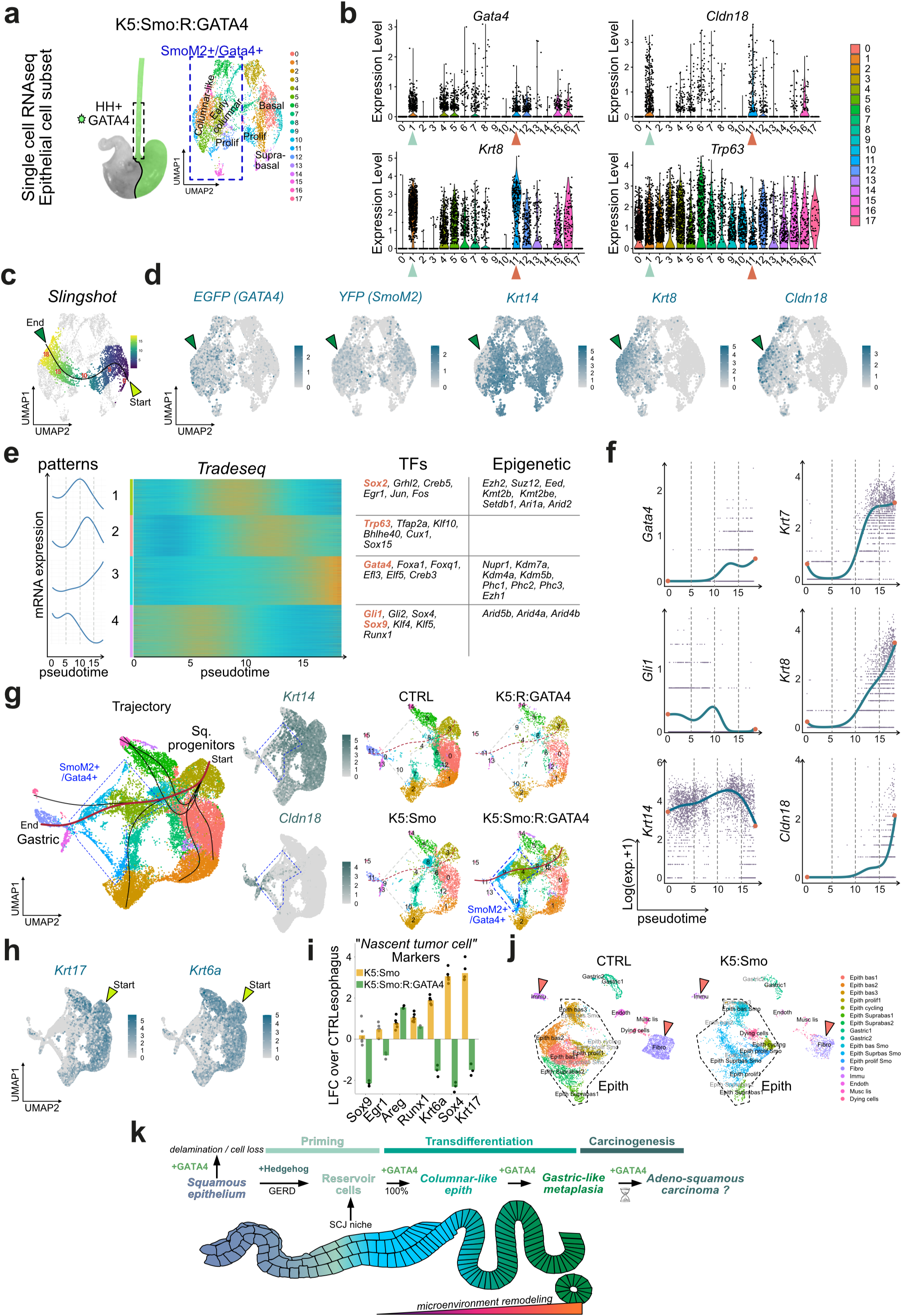
Hedgehog pathway activation induces gastric reprogramming from esophageal progenitors. **a.** Uniform Manifold Approximation and Projection (UMAP) showing cell-type annotation of the epithelial cell subset from the esophagus of K5:Smo:R:GATA4 mice. **b.** Violin plot showing expression of *Gata4*, *Cldn18*, *Krt8* and *Trp63* in the clusters shown in (**a**). Arrows point to *Gata4*+ clusters showing high *Cldn18* and *Krt8* expression and moderate (orange) or low (light blue) *Trp63* expression. **c.** Pseudotime trajectories measured with *Slingshot* in the UMAP from (**a**). **d.** Feature plots showing *EGFP*, *YFP*, *Krt14*, *Krt8* and *Cldn18* expression in K5:Smo:R:GATA4 mouse esophagi scRNA-seq data from (**a**). **e.** *TradeSeq* analysis of dynamic mRNA expression along the pseudotime trajectory illustrated in (**c**), identifying four distinct expression patterns (left). Transcription factors and epigenetic regulators associated with each pattern are shown (right). **f.** Dynamic expression of selected transcripts along the pseudotime trajectory illustrated in (**c**). **g.** Pseudotime trajectories in integrated scRNA-seq data from esophagi of wild-type (CTRL), K5:R:GATA4, K5:Smo, and K5:Smo:R:GATA4 mice. Feature plots of *Krt14* and *Cldn18* expression illustrate the transcriptional proximity of SmoM2+/GATA4+ cells to gastric epithelial cells (cluster 11). **h.** Feature plots showing expression of *Krt17* and *Krt6a* in integrated scRNA-seq data from esophagi of wild-type (CTRL), K5:R:GATA4, K5:Smo, and K5:Smo:R:GATA4 mice. Yellow arrows point to the origin of the transcriptional trajectory shown in (g). **i.** Bar plot showing expression of esophageal nascent tumor cell markers in Smo+ cells from K5:Smo mice and in Smo+/GATA4+ cells from K5:Smo:R:GATA4 mice. **j.** UMAP showing cell-type annotation of merged datasets from the esophagus of CTRL and K5:Smo mice. **k.** Model summarizing our observations

Transcriptomic trajectory analysis using *Slingshot* inferred a differentiation path from basal squamous progenitors toward columnar-like cells (*Krt8*+/Cldn18+/*Krt14*−), consistent with the lineage progression observed *in vivo* by fate mapping (**Fig. 5c, d**). Applying *TradeSeq* to the *Slingshot*-predicted transcriptomic trajectories revealed a dynamic program of transcription factor and epigenetic regulator expression associated with squamous-to-columnar conversion, characterized by a progressive loss of *Sox2*, *Klf4*, *Klf5* and *Trp63* upon co-expression of SmoM2 and GATA4 (**Fig. 5e**). Repression of *Sox2* by GATA4 is consistent with its physiological role during the squamocolumnar junction specification^20^. Notably, the HH target genes *Gli1* and *Gli2* were downregulated as *Gata4* mRNA levels increased, suggesting that GATA4 acts as a negative regulator of HH signaling, as previously reported^29,30^ (**Fig. 5e, f**). Analysis of selected genes confirmed a sequential pattern in which *Krt7* and *Krt8* are upregulated alongside *Gli1* in esophageal cells; subsequent *GATA4* upregulation further increases *Krt7 and Krt8* expression and triggers *Cldn18* expression while driving *Gli1* and *Gli2* downregulation. This sequence suggests that HH signaling needs to be repressed to allow specialized columnar differentiation (**Fig. 5e, f)**.

To further characterize this reprogramming process, we profiled squamous cells from the esophagus as well as the SCJ (LC), from wild-type controls (CTRL), K5:Smo and K5:R:GATA4 mice at single-cell resolution and combined these data with those from K5:Smo:R:GATA4 mice (**Extended Data Fig. 10a, b**). In K5:R:GATA4 samples, only a small number of GATA4-positive cells were detected, consistent with the previously observed elimination of GATA4-expressing cells in the absence of HH activation in the esophagus (**Extended Data Fig. 10a**). Following data integration and cluster annotation, we identified populations of basal progenitors and squamous differentiated cells, as well as distinct GATA4-positive clusters (**Extended Data Fig. 10b-d**).

Further analysis of integrated data revealed a transcriptomic trajectory unique to K5:Smo:R:GATA4 samples, predicting transition of squamous progenitors into GATA4-positive columnar-like cells (**Fig. 5g**). In UMAP representations, this GATA4-positive columnar-like cluster was positioned adjacent to the gastric epithelial cluster, whereas squamous cells from CTRL and K5:Smo samples were located at greater transcriptomic distance (**Fig. 5g**). Examining the origin of this trajectory, we found that the progenitor cells competent for transdfifferentiation were characterized by expression of *Krt17* and *Krt6a* (**Fig. 5h**), as well as *Sox9*, *Areg*, *Egr1* and *Runx1* (**Extended Data Fig. 10e**), which are all transcripts recently associated with nascent esophageal tumors^28^.

According to our data, only esophageal keratinocytes in which HH signaling has been reactivated are permissive to GATA4-driven transdifferentiation. We therefore investigated whether this cell state was directly induced by the HH pathway activation. In line with this hypothesis, immunostaining confirmed the preferential expression of Krt17 in Smo-YFP+ clones in the esophagus (**Extended Data Fig. 11a**). Consistently, bulk RNAseq analysis from samples we generated in a previous work^15^ showed that SmoM2-positive keratinocytes exhibited strong upregulation of several markers associated with tumor initiation in the esophagus (*Sox9*, *Egr1, Areg*, *Runx1*, *Krt6a*, *Sox4* and *Krt17*), which were subequently downregulated upon GATA4 expression (**Fig. 5i**). Interestingly, HH associated genes (*Smo*, *Gli1*, *Gli2*, *Ptch1* and *Ptch2*) were also downregulated except for *Shh* that was even more expressed in SmoM2+/GATA4+ cells than in SmoM2+ cells without GATA4 overexpression (**Extended Data Fig. 11b**).

By merging scRNA-seq data from CTRL and K5:Smo or K5:Smo:R:GATA4 samples, we found that HH pathway activation alone had a limited impact on the microenvironment, while altering epithelial cell states (**Fig. 5j and Extended Data Fig. 11c-e**). By contrast, combined HH activation and GATA4 expression induced more extensive epithelial reprogramming (**Extended Data Fig. 12a-c**), accompanied by pronounced remodeling of the surrounding niche, including the emergence of pro-inflammatory fibroblasts (**Extended Data Fig. 12d**), activation of macrophages, and recruitment of neutrophils, mirroring changes observed at the lesser curvature of K5:Smo:R:GATA4 forestomachs (**Extended Data Fig. 13a-c**). These microenvironmental modifications were associated with increased EGF and CXCL signaling pathways, as observed at the gastric lesser curvature (**Extended Data Fig. 13d-f**).

Taken together, these data support a sequential model in which HH signaling modifies the properties of esophageal progenitors, rendering them permissive to transdifferentiation (“reservoir cells”)^17^, while subsequent GATA4 expression drives columnar conversion and establishes a specialized microenvironment reminiscent of human esophageal metaplasia (**Fig. 5k**).

## Discussion

### Cell plasticity enables oncogenic competence

Current theories of cancer origins diverge in the relative weight they assign to intrinsic genetic alterations and extrinsic tissue-remodelling cues, but converge on the notion that cell state is a key determinant of oncogenic susceptibility^4^. Our findings provide a concrete mechanistic illustration of this principle. Certain oncogenic events require a permissive cell state, induced by injury-driven dedifferentiation, to exert their full transforming potential, and are inert, or even toxic, in the absence of this prior state transition. This model implies that microenvironmental signals, such as those elicited by chronic gastroesophageal reflux, can unlock oncogenic competence by creating a permissive state. In this line, recent work on esophagel squamous cell carcinoma point to a specific phenotype of persister cells that are competent to initiate tumors and create their own niche^28^. Some markers expressed by these persister cells are also upregulated in dedifferentiated foregut epithelium following HH activation. From this state, cells are maintained in the epithelium and seem to become competent to form metaplasia and even neoplasm.

Interestingly, in the lung, three independent studies highlighted that alveolar injury triggers the emergence of a transitional progenitor population, that is absent from homeostatic tissue and arises specifically in response to damage^31–33^. The persistence of this state under chronic injury correlates with disease progression, and a similar transitional progenitor population have been identified within human lung adenocarcinoma tumours^31^. Although the triggering signals may differ (inflammatory cytokines in the lung and HH reactivation in the oesophagus) the underlying logic is conserved: chronic injury sustains a reparative transitional state that, when unresolved, creates conditions permissive for (pre)malignant transformation. Interestingly, markers such as Krt17, Sox9 and Sox4 are shared by these two transitional populations. Analogous precancerous tissue remodelling in response to chronic injury is observed in the pancreas (acinar-to-ductal metaplasia) and cervix (transformation zone)^13^, suggesting that this mechanism may be broadly conserved.

Our findings connect to the broader concept of field cancerization, whereby chronic environmental insults generate a wide zone of molecularly altered, histologically normal epithelium competent for neoplastic transformation^34,35^. Our data suggest that dedifferentiation to an embryonic-like state may represent the cellular basis of such fields in the foregut. Rather than requiring discrete mutational events across many cells, a propagating signal, such as HH pathway induced by gastro-esophageal reflux and/or early metaplastic cells, could prime an expanding epithelial territory for oncogenic reprogramming. Notably, in both human esophageal columnar metaplasia^16^ and our SmoM2/GATA4 model, metaplastic cells themselves produce HH ligands, a finding that raises the possibility that dedifferentiation can propagate from metaplastic cells to neighboring keratinocytes, contributing to the progressive replacement of squamous mucosa. Whether this represents a driver or a consequence of metaplastic spread remains to be determined.

### Embryonic reprogramming as a precancerous reservoir

HH signalling, activated during foregut embryogenesis and silenced in adult differentiated squamous epithelium, is reactivated by chronic reflux and constitutively active in SCJ progenitors^15^. We have shown that this reactivation induces an embryonic-like transcriptomic and epigenetic program in squamous progenitors, reminiscent of oncofetal reprogramming described in other injury contexts^36–39^. Under acute conditions, such reprogrammed states are transient; under chronic injury, they may persist and serve as a reservoir for malignant transformation^17,28^. Our findings suggest that dedifferentiation is not merely a byproduct of tissue stress, but may represent a functional intermediate that creates conditions permissive for metaplasia. During embryogenesis, foregut progenitors are multipotents and depend on specific transcription factor (TF) expression for fate determination. Our model demonstrates that dedifferentiation upon HH activation is sufficient to induce fate commitment driven by the gastric TF GATA4. We could therefore hypothesize that these dedifferentiated cells could act as reservoir for different types of metaplasia and under the expression of intestinal TFs such as GATA6 or CDX2, could undergo intestinal differentiation. Importantly, GATA4 upregulation represses HH signaling while driving columnar conversion, pointing to a mechanism in which successful transdifferentiation would require repressing the dedifferentiated state. This phenomenon is reminiscent of the sequence observed during embryogenesis, in which downregulation of the HH pathway is associated with terminal differentiation of the foregut^40,41^.

### Cellular origin of Barrett’s esophagus

The cellular origin of Barrett’s esophagus remains debated. While it is unambigous that gastric columnar cells can give rise to metaplasia^8,9^, surgical models of chronic reflux in rodents suggested that keratinocytes could also be competent, although it is unclear what proportion of metaplasia may arise from esophageal cells^42,43^. Experimental evidence supports the view that p63+ / Krt5+ squamous progenitors can give rise to intestinal-type metaplasia^10,11^, and that progenitors at the squamo-columnar junction (SCJ), marked by KRT7^23^, possess enhanced plasticity. Importantly, esophageal keratinocytes fate is non-fixed either, and it can be redirected toward an epidermal lineage when the underlying mesenchyme is modified^44,45^.

Our data are consistent with SCJ progenitors being intrinsically more competent for metaplastic conversion than keratinocytes elsewhere such as those along the esophagus. Their proximity to the gastric niche, and constitutive HH pathway activity, likely contribute to this high plasticity. This framework suggests that the higher frequency of metaplasia at the SCJ in humans may reflect the intrinsic competence of these cells rather than their greater exposure to reflux. Metaplastic transformation at the SCJ would therefore depend on a single hit, whereas more proximal sites would require two sequential events. This model would explain why the probability of metaplasia declines as distance from the SCJ increases.

### Oncogene activity is tissue- and state-dependent

Our results also illustrate the context-specificity of oncogene function. *GATA4* and *GATA6* are frequently amplified (40%) or overexpressed in esophageal adenocarcinoma (eAC)^22^, but virtually absent from esophageal squamous cell carcinoma, suggesting a selective advantage that is contingent on the cellular differentiation state. In line with this hypothesis, a recent study reported that a history of gastroesophageal reflux disease (GERD) positively correlates with *GATA4* and *GATA6* amplifications and expression in eAC^46^. GATA6, similarly enriched in intestinal-type gastric and pancreatic cancers, further exemplifies how lineage identity shapes the oncogenic output of a transcription factor. By contrast, GATA4 overexpression in the gastric columnar epithelium has no apparent effect, despite its amplification in a subset of gastric cancers. While this suggests that GATA4 is not oncogenic in this lineage, we cannot exclude the possibility that, as in the foregut squamous lineage, it become oncogenic only in gastric cells that have acquired a permissive state. More broadly, these observations raise the question of whether other foregut developmental regulators display equivalent context-dependent properties, and should motivate systematic analyses of transcription factor activity across tumour subtypes stratified by cell-of-origin.

### Adenosquamous carcinoma: a unified progenitor model

Adenosquamous carcinoma (ASC) is a rare esophageal subtype defined by the coexistence of squamous and glandular components. Two competing hypotheses (collision of two distinct tumours versus bidirectional differentiation from a common progenitor) have long been debated^24,25^. Our model, in which GATA4 overexpression in HH-dedifferentiated squamous progenitors generates lesions with both squamous and columnar characteristics, provides direct mechanistic support for the latter. A single progenitor population with high plasticity, exposed to appropriate environmental and genetic cues, could plausibly give rise to both lineages. Resolving the cellular trajectories underlying ASC, and Barrett’s metaplasia more broadly, will ultimately require lineage barcoding and spatial omics approaches, although the rarity of transitional cell states in human specimens will pose a significant analytical challenge. Of note, our data show that transdifferentiation is associated with tumor initiation, thus reinforcing the continuum between metaplasia and cancer. In human, it is well established that all eAC arise from columnar metaplasia^9,47^. Our data indicate that when metaplasia arises from keratinocytes, the resulting state appears unstable and potentially predisposed toward tumorigenesis. Understanding why transdifferentiated cells may be prone to tumor initiation could offer broader insight into cancer development, not only in the esophagus, but also in malignancies associated with precancerous metaplastic states, including gastric and pancreatic adenocarcinoma and lung squamous cell carcinoma.

## Conclusion

Our findings show that genetic alterations such as *GATA4* amplification, largely inconsequential in a normal epithelial context, become potently transforming within a primed state. This interplay between cell plasticity and oncogene activity positions the dedifferentiated intermediate not as a passive consequence of tissue damage, but as a critical determinant of malignant potential.

## Methods

Our research complies with all relevant ethical regulations. LA1230406 – projects 844N and 793N from the ULB ethical committee fro the experiments on animals and protocol CE 3344 – MTA Beck_1077_IJB Sept 2021 from the ethical Committee from the Institut Jules Bordet (IJB) for experiments with human material.

### Human data analysis

Human gene expression datasets from normal and Barrett’s esophagus of 3 different studies available on Gene Expression Omnibus (GEO) were downloaded and analyzed using GEO2R and R version 4.3.1 (2023-06-16 ucrt) software. Dataset #1: GSE39491, *n*=161 ^48^; #2: GSE34619, *n*=36 ^49^ and #3: GSE36223, *n*=93 ^50^. For each dataset, normality was assessed using the Shapiro–Wilk test. A t-test was applied to normally distributed data (GSE34619), whereas the Mann–Whitney test was used for non-normally distributed data (GSE39491 and GSE36223).

### Experimental model

The Krt5-CreERT2 knock-in mice (The Jackson Laboratory, Stock#029155, RRID:IMSR_JAX:029155) were crossed with the R26SmoM2-YFP mice (The Jackson Laboratory, Stock#005130, RRID:IMSR_JAX:005130) in order to generate K5-CreERT2:R26SmoM2 (K5:Smo) in a C57/BL6N background.

The transgenic TetO-GATA4-IRES-GFP mice were generated in collaboration with the Transgenesis core facility (Institut de Duve at UCLouvain, Belgium) under the supervision of Dr Younes Achouri. The pTRE-β-globulin-Gata4-3×HA-IRES-GFP-polyA insert was randomly integrated in mouse genome. This insert was obtained by subcloning the coding sequence (CDS) of the murine *GATA4* gene into the pTRE-β-globulin-IRES-GFP-polyA plasmid using SacII and NheI restriction sites. Prior to subcloning, this sequence was modified by replacing the CCGCGG codons (124–129) with CCACGA in order to remove a SacII restriction site in the middle of *GATA4* sequence. Full linearized DNA sequence was obtained using XhoI restriction sites and injected for transgenesis. This full linearized DNA sequence is shown in Supplementary Table S1.

The Krt5-CreERT2 knock-in mice were crossed with Rosa26-CAGs-LSL-rtTA3 knock-in mice (The Jackson Laboratory, Stock#029617, RRID: IMSR_JAX:029617) and the TetO-GATA4-IRES-GFP mice in order to genereate Krt5-CreERT2: Rosa26-CAGs-LSL-rtTA3: TetO-GATA4-IRES-GFP mice (K5:R:GATA4).

The K5-CreERT2:R26SmoM2 (K5:Smo) mice were crossed with the Krt5-CreERT2: Rosa26-CAGs-LSL-rtTA3: TetO-GATA4-IRES-GFP mice (K5:R:GATA4) mice in order to genereate Krt5-CreERT2:R26SmoM2:Rosa26-CAGs-LSL-rtTA3:TetO-GATA4-IRES-GFP mice (K5:Smo:R:GATA4).

Unless otherwise specified, a dose of 5 mg tamoxifen (TAM), resuspended in sunflower oil, was administered intraperitoneally to 8-week-old mice to induce recombination of the floxed transgene in the different transgenic mouse models. Following tamoxifen-induced recombination, mice harboring the TetO-GATA4-IRES-GFP allele were fed blue-dyed doxycycline-containing food (1 g/kg) to activate the TetON system. Littermates of the same sex were randomly assigned to experimental groups, and sex was not considered in the experiment design. Mouse colonies were maintained under pathogen-free conditions in a certified animal facility in accordance with the European guidelines, including dark/light cycle, controlled ambient temperature and humidity. All the experiments were approved by the ethical committee from the university (CEBEA at ULB) and conform with regulatory standards (LA1230406 – projects 844N and 793N).

### Method details

For all experiments presented in this study, sample size was large enough to measure the effect size. No randomization and no blinding were performed in this study.

### Histology and Immunostaining on O.C.T Embedded Tissues

For immunostaining on frozen sections, tissues were dissected and embedded in O.C.T. (Tissue Tek, Table S2) and flash frozen for cryopreservation. All the images shown in the figures come from mouse esophagi or forestomach as indicated in the legends and figure labels. For better reproducibility, we focused our histological analyses on the distal segment of the esophagus.

For the following staining: GFP(GATA4), Epcam, Krt7, p63, Cldn18 and Cdx2 tissues were pre-fixed in 4% formaldehyde during 2 h at room temperature, washed in PBS, incubated overnight in a sucrose solution 30% (Table S2) at 4°C, embedded in O.C.T. (Tissue Tek) and flash frozen for cryopreservation.

Samples were sectioned at 6 µm thickness using a M1860 cryostat (Leica Microsystems GmbH). Nonspecific antibody binding was blocked with 5% horse serum (HS), 1% Bovine Serum Albumin (BSA), and 0.2% Triton X-100 during 1 h at room temperature.

Primary antibodies were incubated overnight at 4°C in blocking buffer. Sections were rinsed three times in PBS and incubated for 1 h at room temperature with secondary antibodies in blocking buffer supplemented with Hoechst (4 uM) to stain nuclei. Sections were rinsed three times in PBS. Slides were mounted using Glycergel (Dako) supplemented with 2.5% DABCO (Sigma-Aldrich). The following primary antibodies were used: anti-Krt14 (polyclonal chicken, Biolegend, RRID:AB_2616962), anti-GFP (polyclonal goat, Abcam, RRID:AB_305643), anti-p63 (rabbit monoclonal, Abcam, RRI-D:AB_10971840), anti-Epcam (rat polyclonal, Biolegend, RRID:AB_1089027), anti-Krt7 (rabbit monoclonal, Abcam, RRID:AB_2783822), anti-Krt8 (Rat monoclonal, DSHB, RRID:AB_531826), Cldn18 (rabbit monoclonal, abcam, ab203563), anti-Cdx2 (rabbit monoclonal, abcam, ab76541), anti-Krt17 (rabbit monoclonal, Cell Signaling technology, 4543). Antibodies dilutions are provided in Table S2.

The following secondary antibodies were used: anti-rabbit, anti-goat, anti-rat, anti-chicken, conjugated to AlexaFluor488 (1:500, Jackson ImmunoResearch), to rhodamine Red-X (1:500, Jackson ImmunoResearch) or to Cy5 (1:1000, Jackson ImmunoResearch). Hematoxylin and eosin (H&E) staining was performed following an adapted version of the IHC World protocol. Slides were first fixed for 10 minutes in 4% formaldehyde and then rinsed three times in PBS. Sections were incubated for 45 seconds in Mayer’s hematoxylin (VWR, 10047005) and subsequently rinsed under running tap water for 2 minutes. After a 15-second incubation in 95% ethanol, slides were stained for 30 seconds in eosin (VWR, 10047001). Slides were then progressively dehydrated through one quick bath of 95% ethanol, followed by two 5-minute baths in 100% ethanol and two 5-minute baths in Safe Solvent. Finally, slides were mounted using Safemount. For alcian blue and Periodic Acid –Schiff (PAS) coloration, the NovaUltra Alcian Blue- PAS stain Kit (IHC world, cat# IW-3019) was used following manufacturer recommendations.

### Image acquisition

All samples from the same experiment were imaged with the same exposure settings. Imaging was performed on a Zeiss Axio Imager M2 fluorescence microscope with a Zeiss Axiocam 503 mono camera for immunofluorescence microscopy using Zen Blue 3.2 (Zeiss) software.

H&E and PAS/Alcian Blue-stained slides were scanned using a Hamamatsu NanoZoomer.

### Metaplasia quantification in the different squamous regions of the foregut

Metaplasia incidence was assessed based on Krt14, Cldn18, GFP staining (Krt14−/Cldn18+/GFP+) as well as cellular morphology. A region was considered positive when at least one metaplastic lesion was identified within the analyzed area. In K5:Smo:R:GATA4 mice, a total of 20 greater curvatures, 21 lesser curvatures and 20 esophagus were analyzed. In K5:R:GATA4 mice, 7 samples were analyzed for each region, including the greater curvatures, the lesser curvatures and esophagus.

### Tissue digestion

Murine esophagi and portions of the greater or lesser curvatures of the forestomach were dissected, minced and digested in 2 mg/mL of collagenase I (A&E scientific) for 1h, EDTA 5 mM for 20 min and trypsin 0.125% (A&E scientific) for 10 min. Incubations were performed on a rocking plate at 37°C.

For all the tissues, cells were then rinsed in PBS supplemented with 2% FBS and filtered through a 70 µm cell strainer. All reagents are listed in Table S2.

### FACS isolation

Immunostaining was performed on single cell suspension using PE-conjugated anti-CD45 (1:500, BioLegend), PE-conjugated anti-CD31 (1:500, BioLegend), PE-conjugated anti-CD140a (1:500, BioLegend) and APC-Cy7-conjugated anti-Epcam (clone G8.8, 1:250, Biolegend), for 45 min at 4 °C on a rocking plate. Living epithelial cells were selected by forward scatter, side scatter, doublets discrimination and by Hoechst dye exclusion. Epcam+/Lin- cells were selected based on the expression of Epcam and the exclusion of CD45, CD31, CD140a (Lin-). Fluorescence-activated cell sorting analysis was performed using FACS Aria III and FACS-Diva software (BD Biosciences). For bulk RNA-sequencing, we isolated Epcam High GFP+ cells from the K5:Smo:R:GATA4 condition and total Epcam+ epithelial cells for the control condition (CTRL), as detailed in our previous publication (Vercauteren Drubbel et al. 2021). Used antibodies are listed in Table S2.

### RNA extraction

FACS-sorted cells were collected into TRK lysis buffer (Omega bio-tek) and RNA was extracted using E.Z.N.A Total RNA Kit (Omega bio-tek) according to the manufacturer’s recommendations with DNase I digestion protocol on column (Omega bio-tek).

### RNA-seq and analysis of bulk samples

After RNA extraction, RNA quality was checked using a Bioanalyzer 2100 (Agilent technologies). Indexed cDNA libraries were obtained using the Ovation Solo RNA-Seq System (NuGen) following manufacturer’s recommendations. The multiplexed libraries were loaded on a NovaSeq 6000 (Illumina) using a S2 flow cell and sequencing was performed using a 200 Cycle Kit. Paired-end reads were mapped against the mouse reference genome GRCm38 using STAR software (version 2.5.3a) to generate read alignments for each sample. Gene annotations (Mus_musculus.GRCm38.90.gtf) were obtained from Ensembl (ftp.Ensembl.org). After transcripts assembling, gene level counts were obtained using HTSeq^51^.

Total raw counts were used for subsequent analyzes with degust 4.1.183^52^. All analyses were performed using EdgeR, CPM normalization and ‘‘Min gene read count’’ set at 10, *P*-values were calculated using Quasi-Likelihood (QL) F-test and adjusted (*FDR*) for multiple testing using Benjamini-Hochberg (BH) procedure Bulk RNA-seq was performed on wild-type control mice (CTRL, *n*=4), K5:Smo:R:GATA4 mice treated with doxycycline for 2 weeks starting 2 weeks after tamoxifen injection (HH 2w + GATA4 2w, *n*=3), K5:Smo:R:GATA4 mice treated with doxycycline for 2 weeks starting 6 weeks after tamoxifen injection (HH 6w + GATA4 2w, *n*=4) and K5:Smo:R:GATA4 mice treated with doxycycline for 6 weeks starting 6 weeks after tamoxifen injection (HH 6w + GATA4 6w, *n*=3).

Multidimensional Scaling (MDS) plots were generated using Degust 4.1.183.

Gene expression in GFP+ cells was compared with that in CTRL cells. Genes with an absolute log2 fold change (|*LFC*|) > 2 and a false discovery rate (*FDR*) < 0.05 were considered significantly deregulated. Gastric, intestinal, columnar SCJ, and embryonic esophagus signatures (Extended Data Fig.8) were defined in a previous study as genes upregulated in these columnar tissues compared with wild-type control mice^15^.

The Venn diagrams in Fig.4 and Extended Data Fig.8 were generated using Venny 2.1 by comparing the significantly upregulated transcripts across the different conditions.

### GSEA analysis

Gene set enrichment analysis (GSEA) was performed using the fgsea package^53^ in R version 4.3.1. Genes were ranked in descending order and then filtered to keep only significant differentially expressed genes (*FDR*<0.05).

The ranking metric was defined as the values of LFC. For analysis showed in Fig.4 and Extended Data Fig.8, the C5 collection adapted for mouse which contains gene sets annotated by Gene Ontology (GO) terms was downloaded (http://bioinf.wehi.edu.au/software/MSigDB/). Unbiased enrichment analyses were performed using the “fgseaMultilevel” function. *P*-values were estimated by a two-sided permutation test, with P-values adjusted for multiple testing. Dotplot were generated using “ggplot2” 3.4.2 function in R version 4.3.1.

### Single cell-RNA sequencing and analysis

For esophageal samples (Fig.5 and Extended Data Fig.10, 11, 12, 13) K5:Smo:R:GATA4 and K5:R:GATA4 mice were treated with doxycycline for 6 weeks, starting 6 weeks after tamoxifen (TAM) injection (HH 6w + GATA4 6w). K5:Smo mice were injected with TAM and harvested 12 weeks post-injection, while control (CTRL) mice consisted of untreated wild-type animals. Sorted cells were collected in 1 mL of PBS supplemented with 0.04% BSA and maintained at 4°C. Cells were centrifuged at 300 × g for 10 min at 4°C, resuspended in 50 µL PBS, and cell concentration was determined using a Luna Cell Counter (Westburg Life Sciences). The appropriate amount of cells was subsequently load onto a Chromium chip, to achieve a target recovery of 5,000 to 10,000 cells. Samples were processed using the Chromium Single Cell 3′ Reagent Kits v3.1 (10x Genomics) following the manufacturer’s recommendations. For forestomach samples (Fig.3 and Extended Data Fig.6, 7), the lesser and greater curvatures were microdissected from K5:Smo:R:GATA4 mice treated with doxycycline for 14 weeks, starting 1 week after tamoxifen (TAM) injection (HH 1w + GATA4 14w), and from untreated wild-type control animals (CTRL). Sorted cells were collected in 9 mL of PBS supplemented with 0.04% BSA and mainatained at 4°C. Cells were centrifuged at 300 × g for 10 min at 4°C, resuspended in 50 µL PBS, and cell concentration was determined using a Luna Cell Counter (Westburg Life Sciences). The appropriate amount of cells was subsequently loaded onto a Chromium chip, to achieve a targeted recovery of 5,000 cells. Samples were processed using the GEM-X multiplex v4.1 kit (10x Genomics), following the manufacturer’s recommendations.

The multiplexed libraries were loaded on a NovaSeq 6000 (Illumina) using a S1 flow cell and paired-end sequences were produced using a 100-cycle kit (Read1, 28 cycles; i7 Index, 8 cycles; i5 Index, 0 cycles; and Read2, 87 cycles). The Cell Ranger pipeline (v3.1.0), incorporating the STAR RNA-seq aligner, was used to generate output files aligned to the mm10 reference genome, with custom EYFP and GFP sequences added (mCherry was included but not used in this analysis). Package Seurat v5.3 from Bioconductor^54^ was used in R v4.4.2 to perform all the analyses.

The analyses followed recommendations from Satija Lab (https://satijalab.org). The datasets were converted into Seurat objects and the number of sequenced cells is reported in “Nbr of cells before subset” column of Table S3. SoupX v1.6.2^55^ was used with default parameters to remove ambient RNA contamination. For each sample, low-quality cells, empty droplets, and multiplets were excluded based on mitochondrial gene content and nCount_RNA. Sample-specific thresholds are provided in the “Subset” column of Table S3. The data were normalized using SCTransform^56^, with the parameter vars.to.regress = c(“percent.mt”, ”percent.rb”), where percent.mt and percent.rb were computed using PercentageFeatureSet(pattern = "^mt-") and PercentageFeatureSet(pattern = "^Rp[sl]"), respectively, to minimize the influence of mitochondrial and ribosomal reads in downstream analyses. For each Seurat object, the following steps have been run: Principal component analysis (PCA) was performed using RunPCA, with the number of retained principal components (npcs) determined based on the variance explained. Sample-specific npcs are provided in the “npcs” column of Table S3. Uniform Manifold Approximation and Projection (UMAP) analyses have been done using “RunUMAP”. Distances between cells were defined using “FindNeighbors”. Cells were clustered using FindClusters, with a resolution parameter defined for each sample as reported in the “resolution” column of Table S3. Doublets were subsequently removed using the DoubletFinder R package (v2.0.4), and the final number of cells after preprocessing is reported in the “Nbr of cells after subset” column of Table S3. For merged samples (see “Combination” column Table S3), the merge function was used with the parameter merge.data = TRUE, and variable features were reassigned from the original objects using the command “SelectIntegrationFeatures” as recommended. For integrated objects (see “Combination” column Table S3), following satija lab pipeline, merged datasets were integrated using the IntegrateLayers function with the parameter method = RPCAIntegration. The epithelial subset was defined based on the selection of *Epcam*-*Cdh1*-expressing clusters and the exclusion of *Pdgfra*-, *Vim*-, *Ptprc*-, and *Pecam1*-expressing clusters. Excluded clusters are detailed in the “Subset” column of Table S3.

Following subset, merging or integration, PCA, UMAP, FindNeighbors, and FindClusters were then performed using the parameters reported in Table S3. Last, for visualization and differential expression (DE) analysis, RNA counts were normalized using “NormalizeData.” Differentially expressed markers for each group versus all others were identified using the Wilcoxon test in FindMarkers with the following parameters: *logfc.threshold* = 0.2, *min.pct* = 0.1, and *min.diff.pct* = 0.1. Genes with adjusted *P*-values < 0.05 were considered significant. Cluster annotations were performed using the list of significantly up- and downregulated genes in each cluster.

Violin plots and feature plots were generated using the VlnPlot and FeaturePlot functions, respectively. For markers expressed in very few cells, the order parameter was set to TRUE.

Slingshot v2.10.0^57^ was used to perform pseudotime analysis, with starting points selected based on biological knowledge. Gene expression dynamics along pseudotime were explored using the tradeSeq package^58^, following recommended workflows.

Characterization of the signaling pathway network was performed using CellChat v1.6.1^59^, following the authors’ recommendations.

### Statistics and Reproducibility

Statistical and graphical data analyses were performed using R software. For each analysis, the R software version is specified in the corresponding section. All experiments presented were independently replicated at least three times and included a minimum of three biological replicates. Data in histograms represent the mean unless otherwise specified in the legend Statistical tests used for data analysis are stated in the legend. *P*-value < 0.05 was considered statistically significant.

## Acknowledgments

We thank the FACS core facility and the animal house facility of Université libre de Bruxelles (ULB) for their help. Sequencing was performed at the Brussels Interuniversity Genomics High Throughput core (www.brightcore.be) by Dr Frédérick Libert and Anne Lefort. We thank Dr Younes Achouri and Prof Patrick Jacquemin from the Transgenesis core facility (Institut de Duve at UCLouvain, Belgium) for the generation of the TetO-GATA4-IRES-GFP mouse strain.

This work was supported by Fondation contre le Cancer (FCC/ULB 2018-067 and 2022-164) and the the Fonds de la Recherche Scientifique - FNRS under Grant (CDR n°400008574) with BB as Principal Investigator. BB is Senior Research Associate and AVD a Postdoctoral Researcher of the Fonds de la Recherche Scientifique – FNRS at ULB.

## Author contributions

AVD and BB designed, validated and conducted experiments. AVD performed single-cell RNA sequencing data analysis. SP provided technical help. AVD and BB conceived the project, supervised experiments, and wrote the manuscript. Funding acquisition and supervision of the project by BB. Preparation of the manuscript by all authors.

