## Extended data Figure 1-13 and Supplementary table 1-3 for "Dedifferentiation unlocks keratinocyte competence for metaplasia and tumorigenesis in the foregut"

### Extended Data Fig. 1

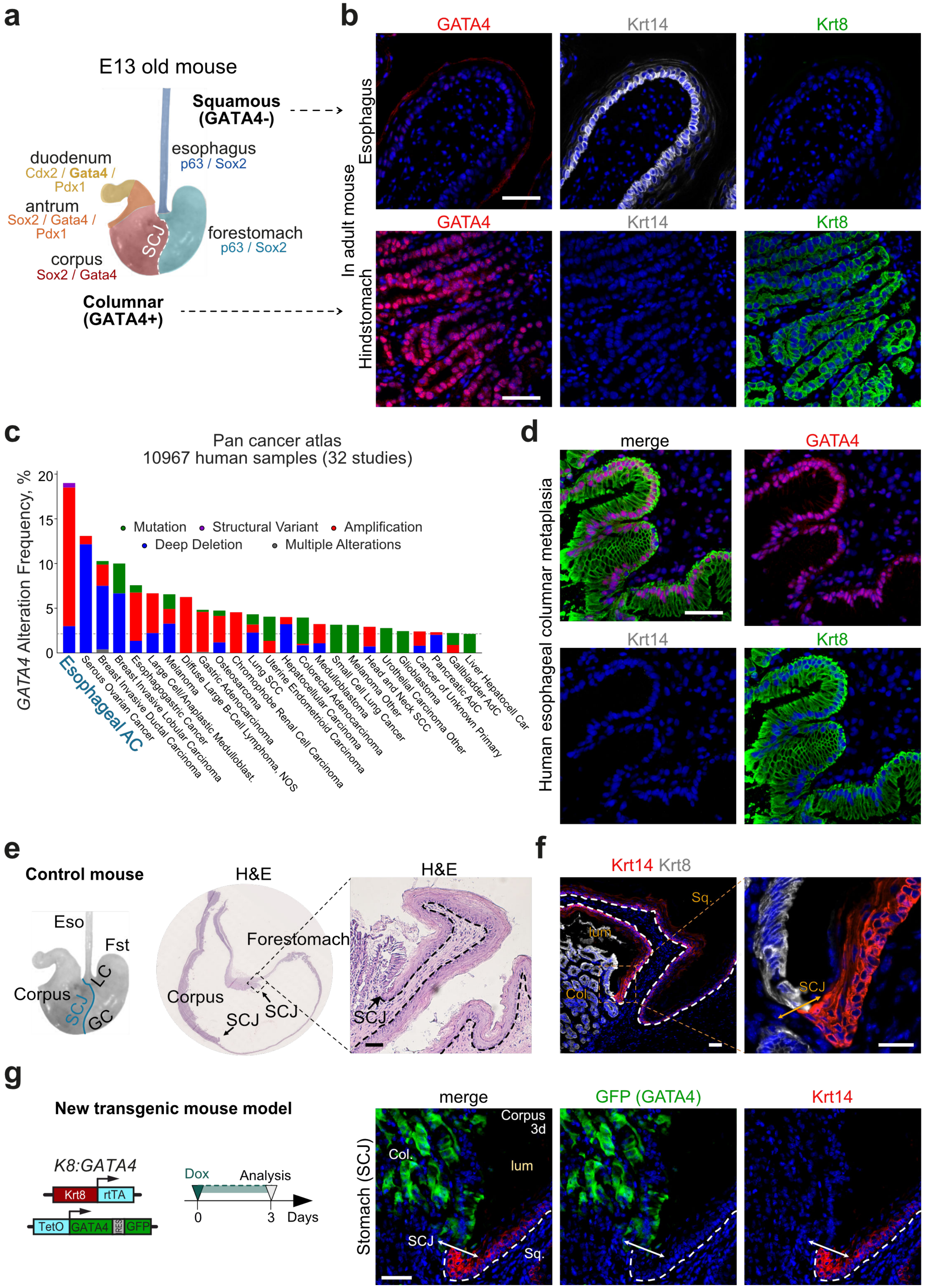

**Extended Data Fig. 1**

**a.** Schematic representation of the pattern of transcription factor expression in embryonic mouse foregut. The scheme is adapted from Willet and Mills[18].

**b.** Immunostaining for GATA4, Krt14, and Krt8 in squamous esophageal epithelium and gastric columnar mucosa.

**c.** Frequency (%) of *GATA4* genomic alterations across 32 human cancer types. Data were filtered to include only cancer types with *GATA4* alteration frequencies >2%.

**d.** Immunostaining for GATA4, Krt14, and Krt8 in human esophageal columnar metaplasia. Merged images are shown in Fig. 1e.

**e.** Hematoxylin and eosin (H&E) staining of mouse stomach showing squamo-columnar junction (SCJ) histology in control wild type mice.

**f.** Immunostaining for Krt14 and Krt8 illustrating the typical keratin expression shift at the SCJ in control wild type mice.

**g.** Immunostaining for GFP and Krt14 at the SCJ in K8-rtTA:GATA4 (K8:R:GATA4) mouse stomach, 3 days after initiation of Dox treatment.

SCJ, squamocolumnar junction; Sq, squamous; Col, columnar; GC, greater curvature; LC, lesser curvature; Fst, forestomach; Eso, esophagus; Dox, Doxycycline.

Nuclei are counterstained with Hoechst (blue) in all images. Scale bar: 50 µm unless otherwise indicated.

### Extended Data Fig. 2

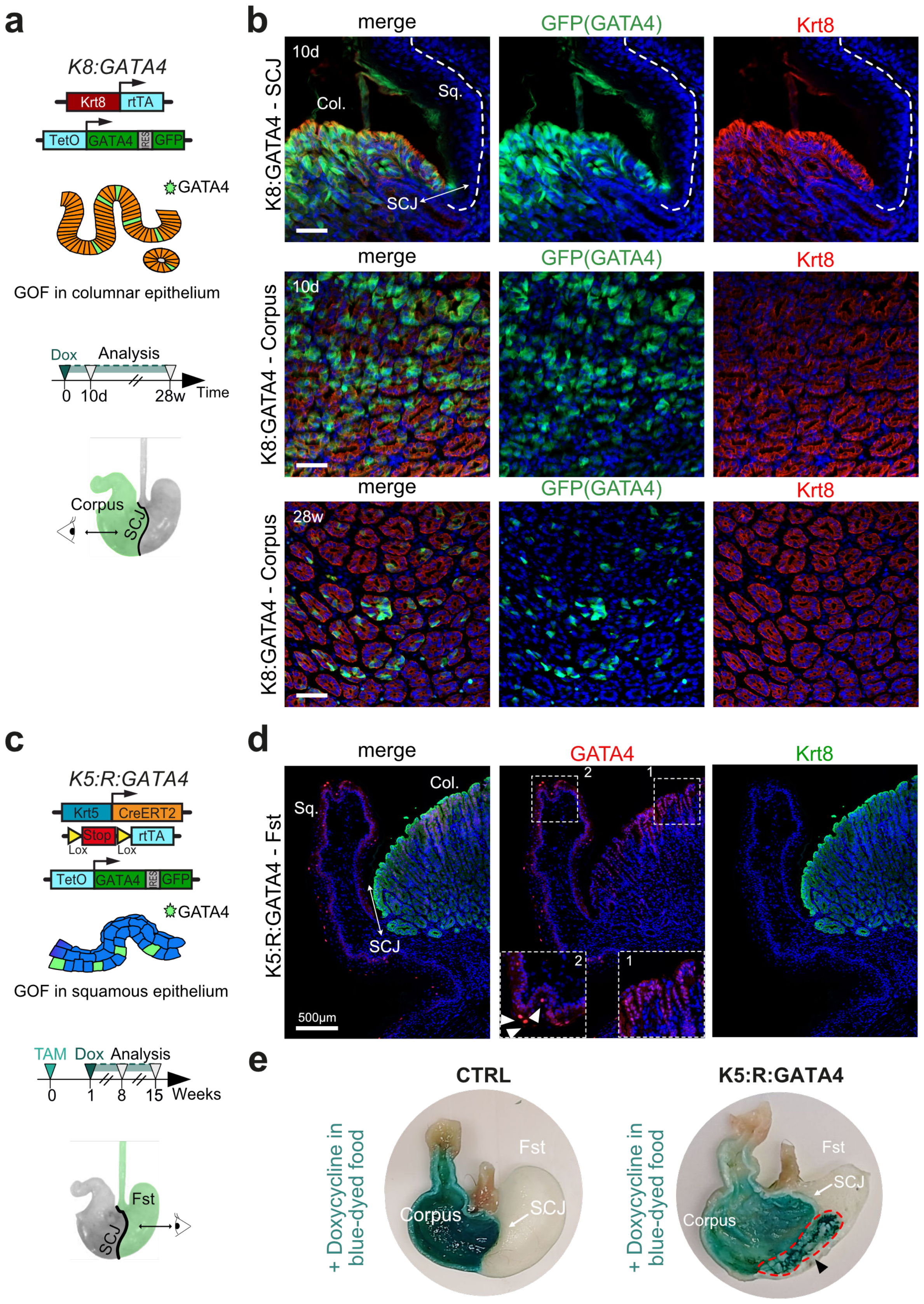

**Extended Data Fig. 2**

**a.** Experimental design to assess the effect of GATA4 overexpression in Krt8-positive gastric columnar epithelium.

**b.** Immunostaining for GFP (ectopic GATA4) and Krt8 in gastric columnar mucosa of K8:GATA4 mice at 10 days and 28 weeks after initiation of Dox treatment. Merged channel images are also shown in Fig. 1g.

**c.** Experimental design to assess the effect of GATA4 overexpression in Krt5-positive squamous epithelium of the foregut.

**d.** Immunostaining for GATA4 and Krt8 in the SCJ of the gastric mucosa in K5CreER:Rosa26-rtTA:TetO-GATA4-GFP (K5:R:GATA4) mice, 7 weeks after Dox treatment initiation. The GATA4 transgene is expressed at a higher level in a subset of squamous cells compared with endogenous GATA4 expression observed in the columnar epithelium.

**e.** Macroscopic image of stomachs from CTRL and K5:R:GATA4 mice after 14 weeks on a blue-dyed Dox diet. Blueish staining of the mucosa confirms Dox exposure but does not reflect GATA4 expression.

GOF, gain of function; d, days; w, weeks; SCJ, squamocolumnar junction; Sq, squamous; Col, columnar; Fst, forestomach; TAM, tamoxifen; Dox, Doxycycline.

Nuclei are counterstained with Hoechst (blue) in all images. Scale bar: 50 μm unless otherwise indicated.

### Extended Data Fig. 3

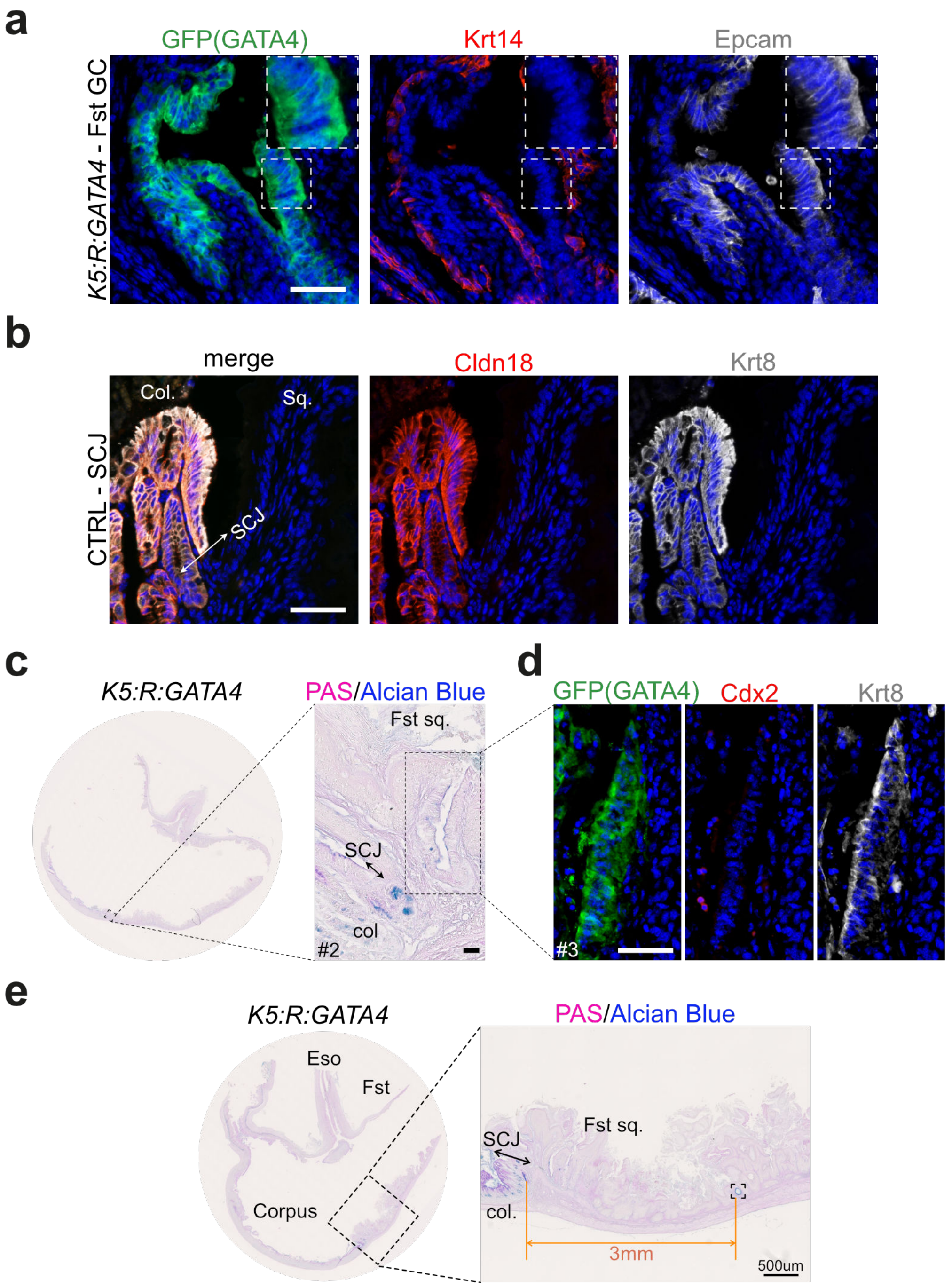

**Extended Data Fig. 3**

**a.** Immunostaining for GATA4, Krt14, and Epcam in the greater curvature (GC) of the forestomach of K5:R:GATA4 mice, 14 weeks after initiation of Dox treatment. Merged channel images are shown in Fig. 1j. GFP labeling identifies keratinocytes overexpressing ectopic GATA4, which show heterogeneous upregulation of Epcam and downregulation of Krt14.

**b.** Immunostaining for Cldn18 and Krt8 at the GC squamo-columnar junction (SCJ) in control (CTRL) animals. Merged channel images are also shown in Fig. 1k. Cldn18 and Krt8 expression is confined to the gastric columnar epithelium and absent from the squamous lineage.

**c.** Periodic acid–Schiff (PAS)/Alcian blue staining with hematoxylin counterstain in K5:R:GATA4 mouse stomach, 14 weeks after initiation of Dox treatment, showing the localisation of the lesion shown in Fig. 1l with mucin-producing cells in the vicinity of the SCJ along the GC.

**d.** Immunostaining of the lesion shown in (**c**) for GATA4, Cdx2 and Krt8 in *de novo* columnar-like epithelium arising in the forestomach of K5:R:GATA4 mice. The absence of Cdx2 expression indicates that GATA4 overexpression does not induce intestinal metaplasia. Panels (**c**) and (**d**) were performed on serial sections.

**e.** Periodic acid–Schiff (PAS)/Alcian blue staining with hematoxylin counterstain in K5:R:GATA4 mouse stomach, showing mucin-producing cells located along the GC, distal to the SCJ. Cropped image is shown in Fig. 1m.

SCJ, squamocolumnar junction; GC, greater curvature; Fst, forestomach; Eso, esophagus; Dox, Doxycycline.

Nuclei are counterstained with Hoechst (blue) in all images. Scale bar: 50 µm unless otherwise indicated.

### Extended Data Fig. 4

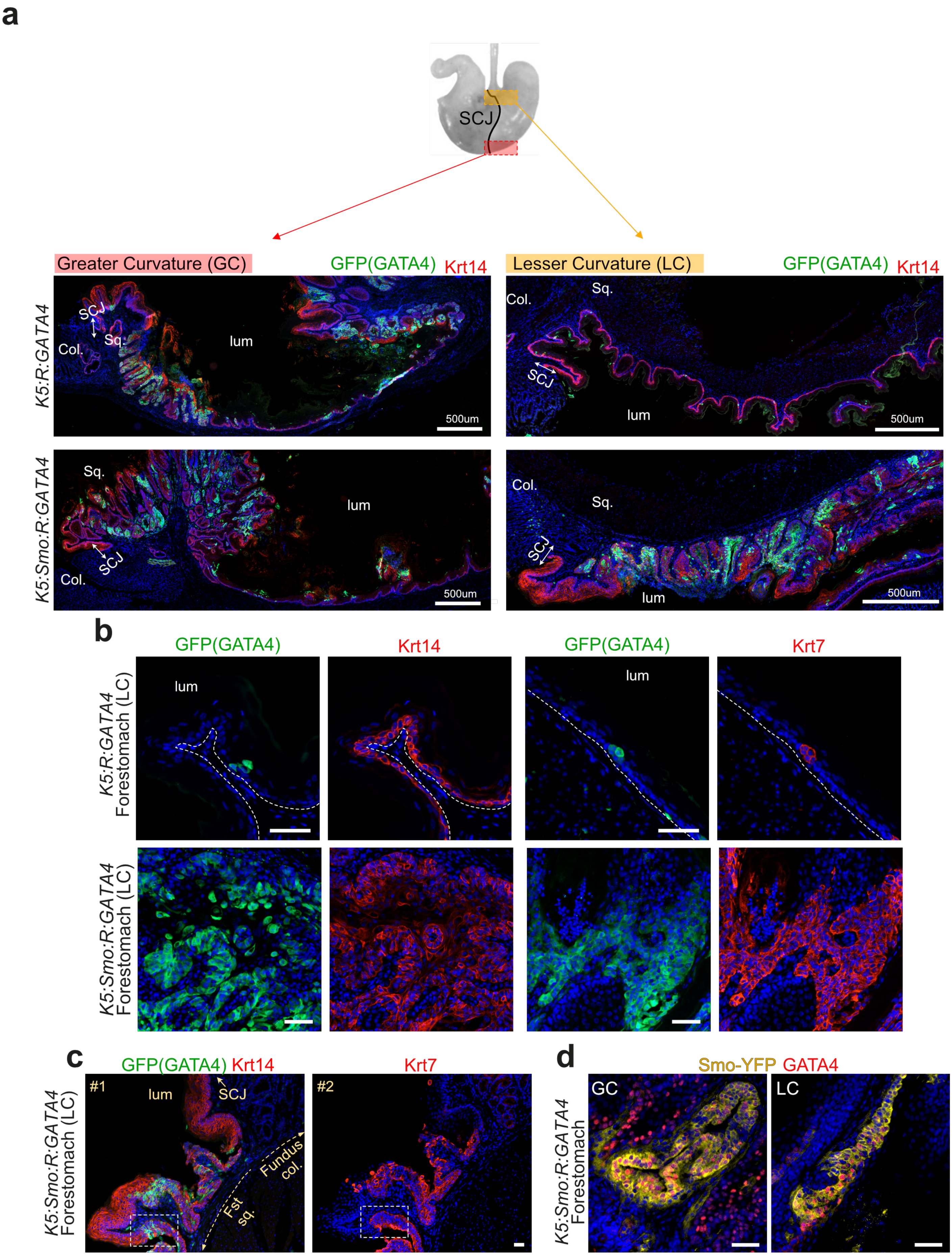

**Extended Data Fig. 4**

**a.** Immunostaining for GFP (ectopic GATA4) and Krt14 in the greater curvature (GC – left pictures) or lesser curvature (LC – right pictures) of the stomach in K5:R:GATA4 (top) and K5:Smo:R:GATA4 (bottom) mice.

**b.** Immunostaining for GFP (ectopic GATA4) and Krt14 or Krt7 in the LC of the stomach in K5:R:GATA4 (top) and K5:Smo:R:GATA4 (bottom) mice.

**c.** Immunostaining for GFP (ectopic GATA4) and Krt14 or Krt7 in the SCJ area of the LC of the stomach in K5:Smo:R:GATA4 mice. Krt14 and Krt7 staining was performed on serial sections. These stainings should be compared with the hematoxylin and eosin (H&E) staining of the corresponding region shown in Fig. 2d.

**d.** Immunostaining for GATA4 and SmoM2-YFP in the GC or LC of the stomach in K5:Smo:R:GATA4 mice.

All mice shown in this figure were fed doxycycline for 14 weeks (GATA4 expression), starting 1 week after tamoxifen induction.

SCJ, squamocolumnar junction; GC, greater curvature; LC, lesser curvature; Col, columnar; Sq, squamous;

Nuclear Hoechst staining is shown in blue in all images. Scale bar: 50 µm, unless otherwise indicated.

### Extended Data Fig. 5

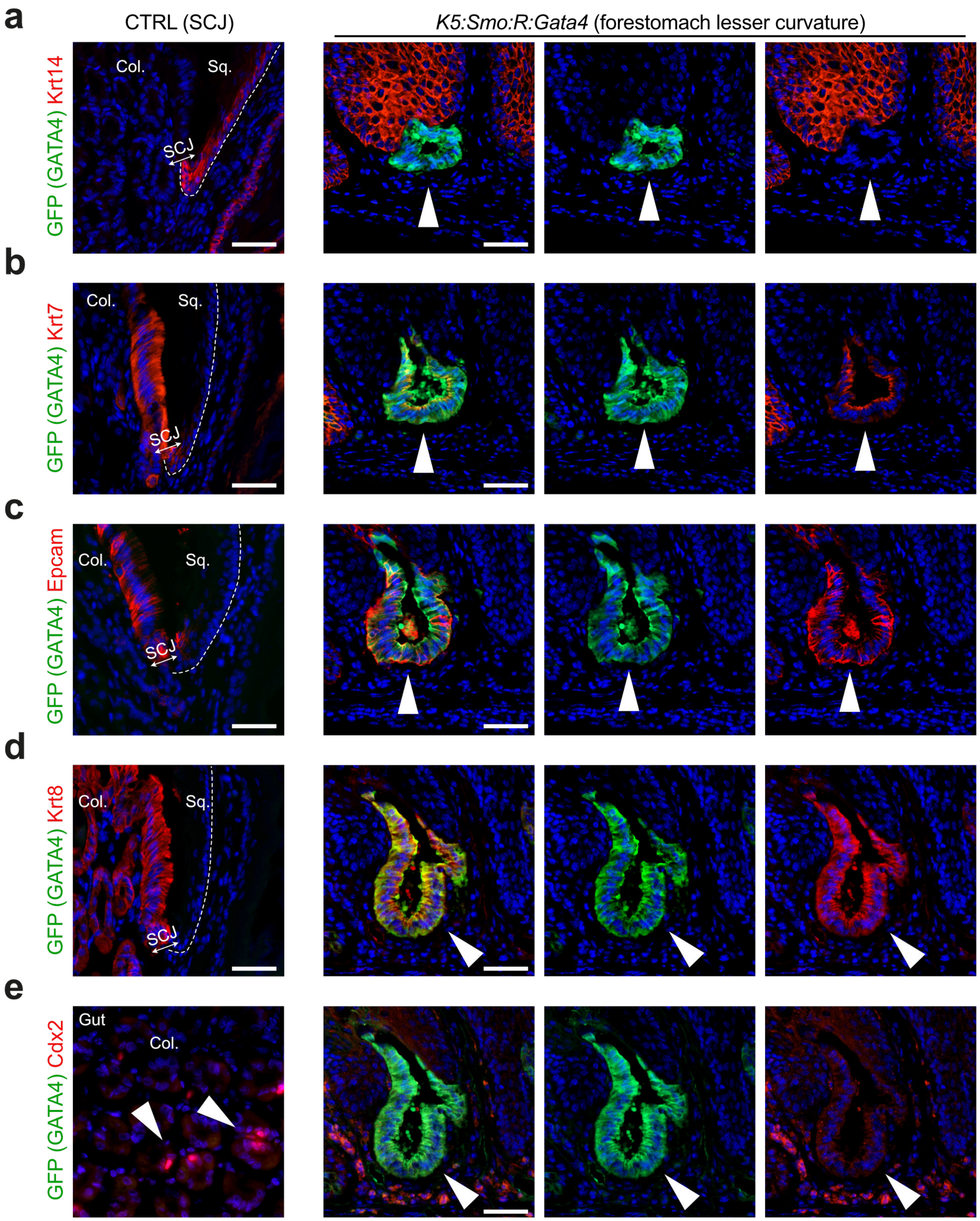

**Extended Data Fig. 5**

**a.** Immunostaining for GFP (ectopic GATA4) and Krt14 in the lesser curvature of the forestomach from K5:Smo:R:GATA4 mice, fed doxycyclin for 14 weeks, showing a metaplastic lesion located away from the SCJ. Staining at the squamo-columnar junction (SCJ) from control (CTRL) animals is shown as a positive control.

**b.** Same as in (a) with GFP (GATA4) and Krt7.

**c.** Same as in (a) with GFP (GATA4) and Epcam.

**d.** Same as in (a) with GFP (GATA4) and Krt8.

**e.** Same as in (a) with GFP (GATA4) and Cdx2. Staining in intestinal mucosa from CTRL animals is shown as a positive control (left image).

Immunostainings for GFP with Krt14 or p63, and for GFP with Cldn18, in the same region are shown in Fig. 1h,i. All panels were performed on serial sections.

SCJ, squamocolumnar junction; Col, columnar; Sq, squamous

Nuclear Hoechst staining is shown in blue in all images. Scale bar: 50 µm.

### Extended Data Fig. 6

**a**

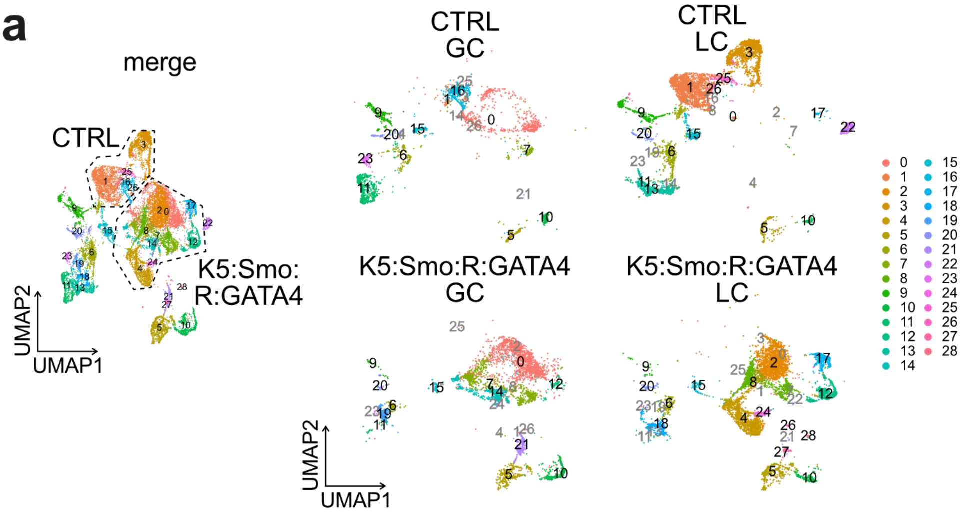

**b**

Number of cells per cluster

| | CTRL | | K5:Smo:R:GATA4 | | |
|---|---|---|---|---|---|
| ID | GC | LC | GC | LC | Annotation |
| 0 | 744 | 26 | 1728 | 17 | Epith |
| 1 | 64 | 2125 | 3 | 15 | Epith prolif |
| 2 | 0 | 5 | 8 | 1803 | Epith |
| 3 | 7 | 1495 | 0 | 17 | Epith suprab |
| 4 | 1 | 3 | 4 | 1339 | Epith |
| 5 | 154 | 164 | 363 | 414 | Immu1 |
| 6 | 137 | 435 | 116 | 202 | Fibro1 |
| 7 | 108 | 5 | 742 | 9 | Epith prolif |
| 8 | 0 | 7 | 6 | 842 | Epith |
| 9 | 262 | 445 | 21 | 34 | Smooth Muscle |
| 10 | 124 | 74 | 362 | 198 | Immu2 |
| 11 | 662 | 44 | 31 | 9 | Fibro2 |
| 12 | 0 | 0 | 79 | 629 | Epith suprab |
| 13 | 0 | 629 | 0 | 23 | Fibro3 |
| 14 | 5 | 2 | 577 | 0 | erythrocyte |
| 15 | 124 | 239 | 70 | 98 | Endoth |
| 16 | 441 | 15 | 0 | 0 | Epith |
| 17 | 0 | 25 | 0 | 409 | Epith differentiated |
| 18 | 0 | 0 | 0 | 338 | Fibro4 |
| 19 | 0 | 1 | 310 | 12 | Fibro5 |
| 20 | 126 | 79 | 38 | 57 | Pericytes |
| 21 | 1 | 0 | 267 | 18 | Immu3 |
| 22 | 0 | 222 | 0 | 5 | Epith prolif |
| 23 | 160 | 4 | 6 | 3 | Fibro6 |
| 24 | 0 | 0 | 7 | 135 | Epith |
| 25 | 4 | 128 | 1 | 2 | Epith suprab |
| 26 | 4 | 91 | 7 | 25 | Epith |
| 27 | 0 | 0 | 0 | 57 | Immu4 |
| 28 | 0 | 0 | 0 | 38 | Immu5 |

**c**

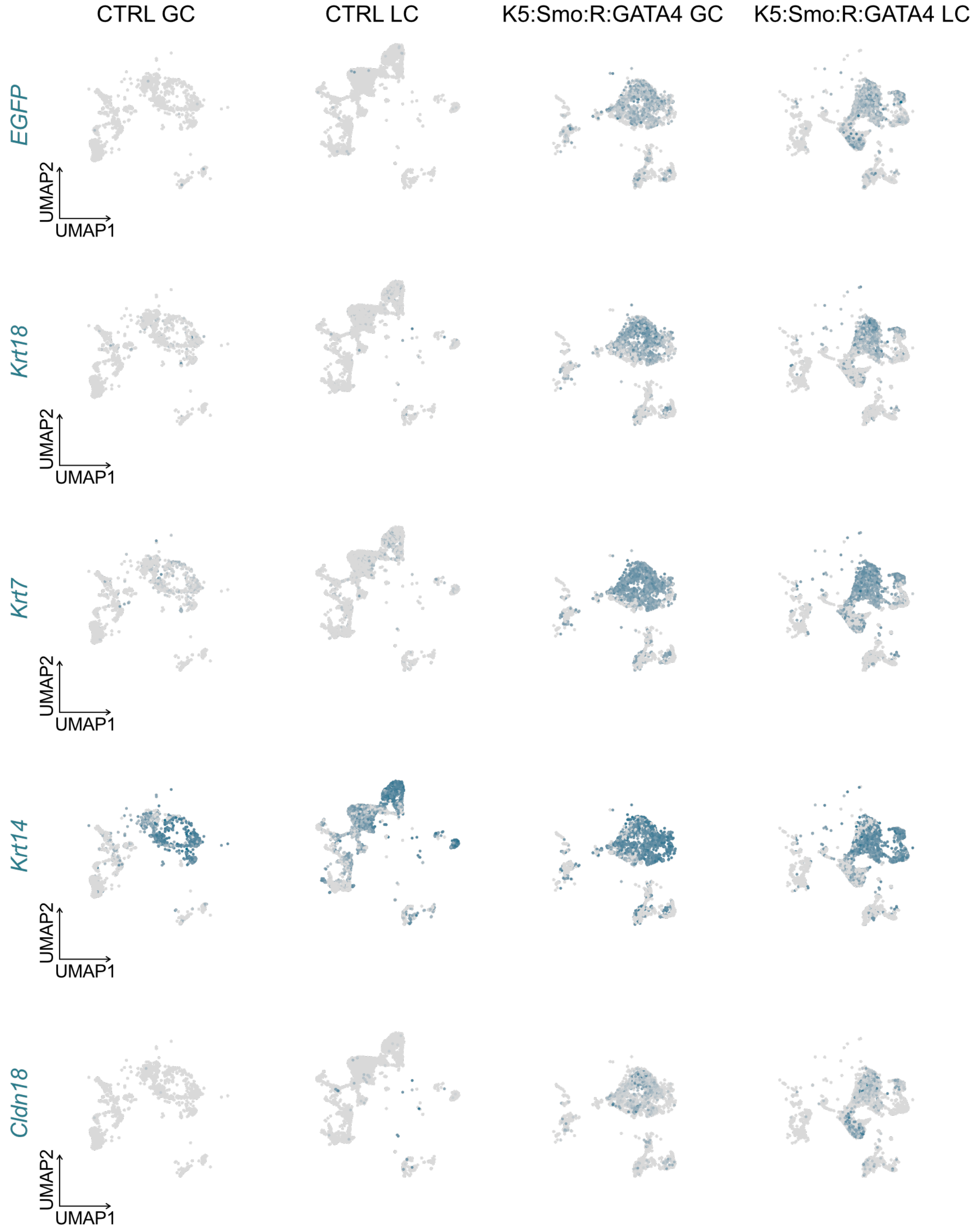

**Extended Data Fig. 6**

**a.** UMAP of merged scRNA-seq data from the greater curvature (GC) and lesser curvature (LC) of control mice (CTRL GC and CTRL LC, respectively), and from K5CreER:Rosa-rtTA:SmoM2:GATA4 mice (K5:Smo:R:GATA4 GC and K5:Smo:R:GATA4 LC, respectively). Right panels show data split by condition.

**b.** Table summarizing the number of cells per cluster and per condition. Values are shown in grey when fewer than 30 cells are present. Mutant conditions are highlighted in light green.

**c.** Feature plots showing the expression of the indicated markers across the four conditions. For markers with low overall expression (*EGFP*, *Cldn18*), feature plots were generated with order = TRUE to improve visualization.

### Extended Data Fig. 7

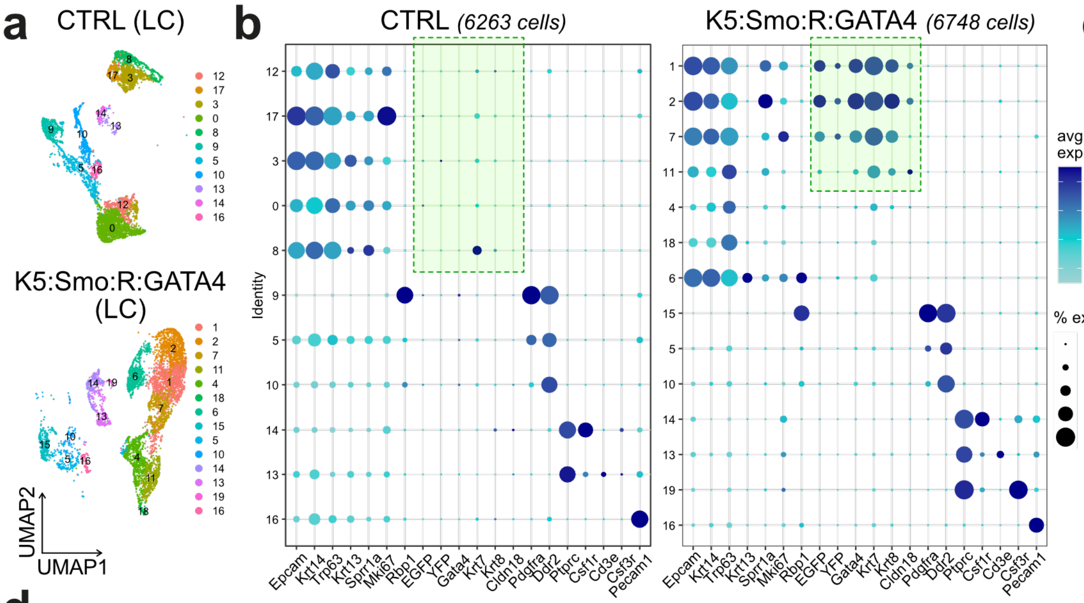

**c** Number of cells per cluster

| ID | CTRL | K5:Smo:R:GATA4 | Tot | Annotation |
|---|---|---|---|---|
| 12 | 502 | 16 | 518 | Bas Prolif1 |
| 17 | 182 | 3 | 185 | Bas Prolif2 |
| 3 | 988 | 13 | 1001 | Bas+Early suprab |
| 0 | 1823 | 13 | 1836 | Bas+Suprab |
| 8 | 590 | 27 | 617 | Suprab1 |
| 9 | 608 | 6 | 614 | Fibro1 |
| 1 | 3 | 1326 | 1329 | SmoGata4-1 |
| 2 | 5 | 998 | 1003 | SmoGata4-2 |
| 7 | 0 | 685 | 685 | SmoGata4-Prolif |
| 11 | 1 | 519 | 520 | Bas commited1 |
| 4 | 1 | 865 | 866 | Bas commited2 |
| 18 | 0 | 97 | 97 | Bas altered |
| 6 | 4 | 696 | 700 | Suprab2 |
| 15 | 0 | 357 | 357 | Fibro2 |
| 19 | 0 | 59 | 59 | Neutrophils |
| 5 | 607 | 249 | 856 | Fibro3 |
| 10 | 516 | 79 | 595 | Muscle |
| 14 | 114 | 287 | 401 | Macrophages |
| 13 | 113 | 360 | 473 | Lymphocytes |
| 16 | 206 | 93 | 299 | Endoth |

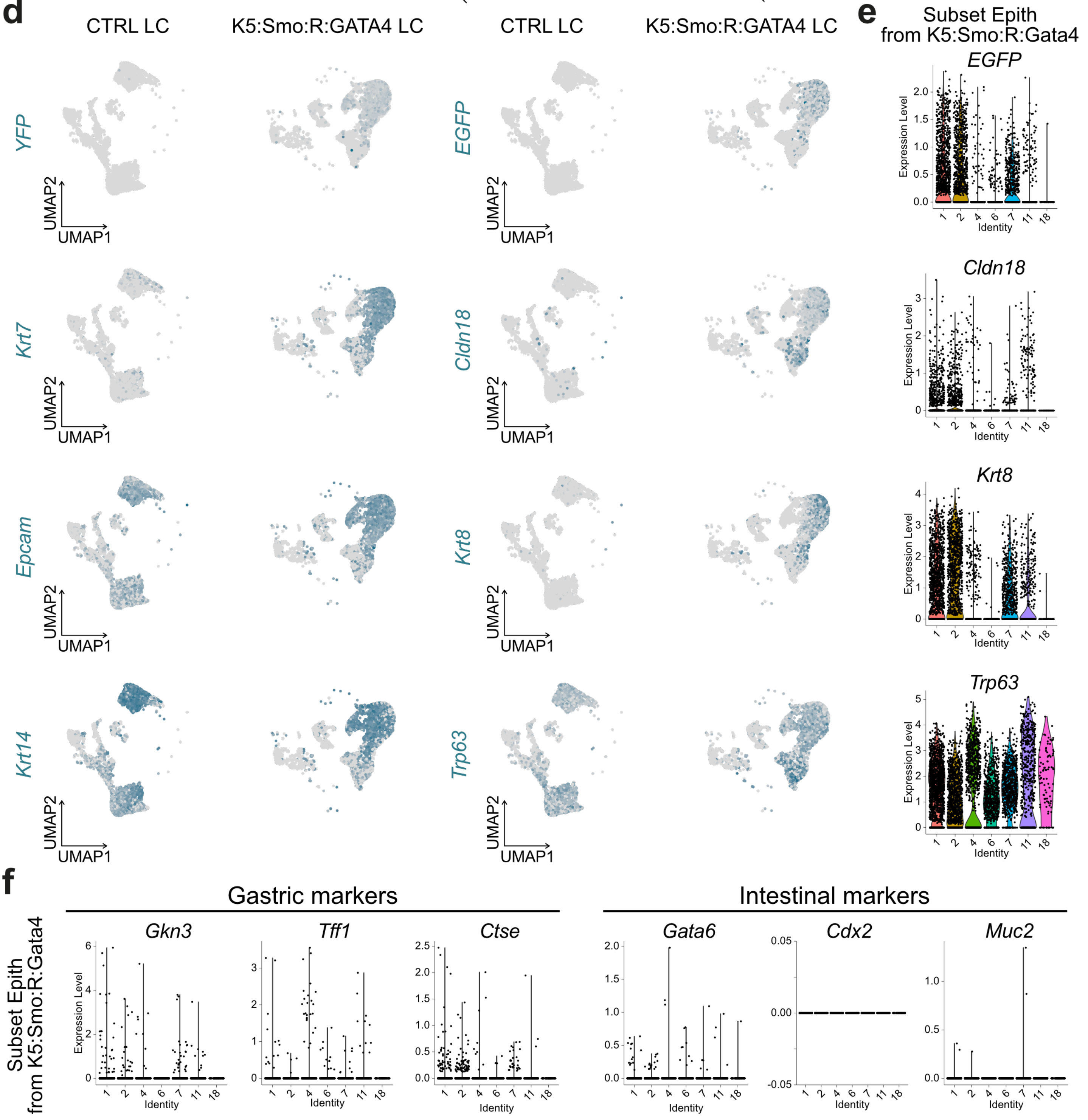

**Extended Data Fig. 7**

**a.** UMAP of scRNA-seq data from lesser curvature (LC) cells of control (CTRL) and K5CreER:Rosa-rtTA:SmoM2:GATA4 (K5:Smo:R:GATA4) mice.

**b.** Dot plot showing the expression of selected marker genes used for cluster annotation in the scRNA-seq dataset from CTRL and K5:Smo:R:GATA4 mice. Dots associated with genes overexpressed in mutant condition are highlighted in green.

**c.** Table summarizing the number of cells per cluster and per condition. Values are shown in grey when fewer than 30 cells are present. Mutant conditions are highlighted in light green.

**d.** Feature plots showing the expression of squamous markers (*Krt14*, *Trp63*), metaplasia-associated markers (*Cldn18*, *Krt7*, *Krt8*, *Epcam*), and lineage reporters (*YFP*, *EGFP*) in scRNA-seq data from CTRL and K5:Smo:R:GATA4 mice. For markers with low overall expression (*EGFP*, *Cldn18*), feature plots were generated with order = TRUE to improve visualization.

**e.** Violin plots showing heterogeneous expression of *EGFP*, *Cldn18*, *Krt8*, and *Trp63* in a subset of epithelial cells from K5:Smo:R:GATA4 mice.

**f.** Violin plots showing expression of gastric markers (*Gkn3*, *Tff1*, *Ctse*) in clusters 1, 2 and 7, and the virtual absence of intestinal markers (*Gata6*, *Cdx2*, *Muc2*) in epithelial cells from K5:Smo:R:GATA4 mice.

### Extended Data Fig. 8

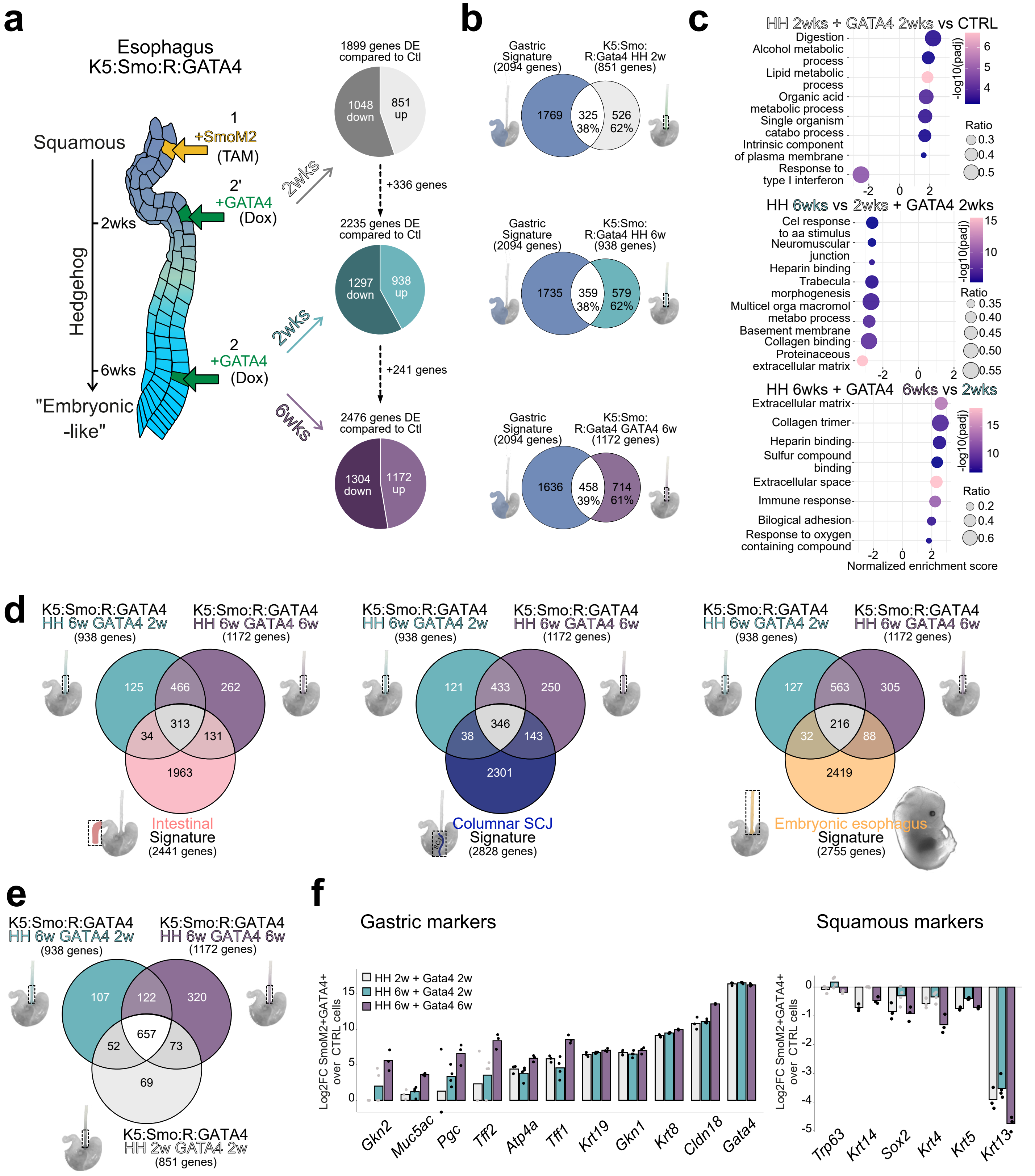

**Extended Data Fig. 8**

**a.** Experimental design with varying intervals between TAM-induced SmoM2 activation and Dox-induced GATA4 expression (left). Pie charts show the number of significantly up- and down-regulated genes relative to control (CTRL) esophageal epithelium ($|LFC| > 2$, $FDR < 0.05$) analyzed following bulk RNAseq of the three time points (right).

**b.** Venn diagram showing the overlap between genes upregulated in GATA4+ cells in the three time point described in (a) and genes upregulated in gastric columnar epithelial cells compared to control esophageal squamous progenitors.

**c.** Gene set enrichment analysis (GSEA) of differentially expressed genes comparing the three time points defined in (a).

**d.** Venn diagrams showing the overlap between transcriptional profiles of esophageal progenitors after 6 weeks of HH pathway activation followed by 2 or 6 additional weeks of GATA4 overexpression, and predefined transcriptomic signatures of intestinal, columnar junction, and embryonic epithelial cells.

**e.** Venn diagrams showing the overlap in gene expression between esophageal progenitors at the three time points defined in (a).

**f.** Bar charts summarizing the relative expression of gastric and squamous markers in GFP+ FACS-sorted cells compared to CTRL squamous esophageal progenitors at the three time points defined in (a). Black dots indicate genes with $FDR < 0.05$ relative to CTRL.

Wks, weeks; Hedgehog, HH; Dox, Doxycycline; TAM, tamoxifen.

### Extended Data Fig. 9

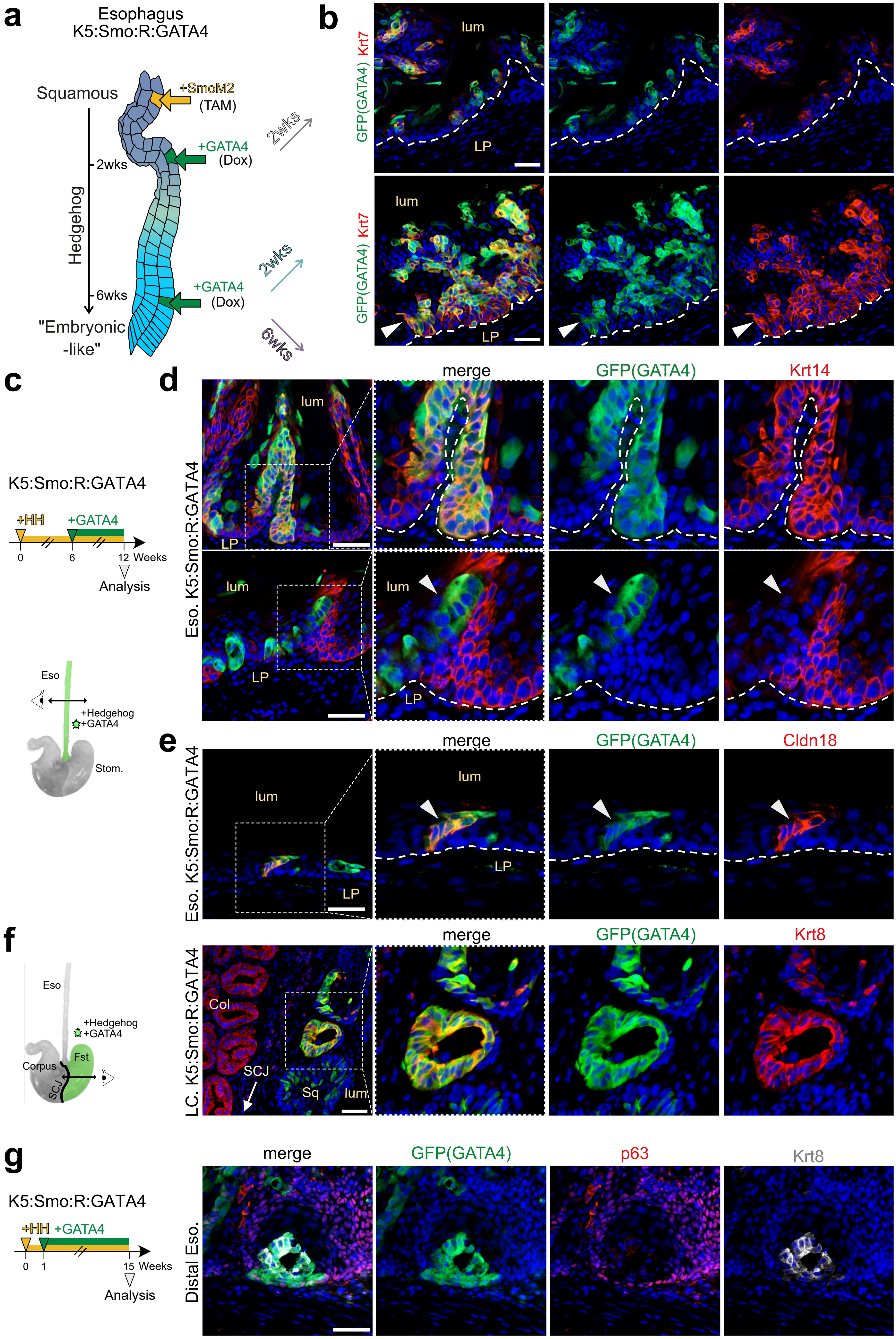

**Extended Data Fig. 9**

**a.** Experimental design with varying intervals between tamoxifen (TAM)-induced SmoM2 activation and Doxycycline (Dox)-induced GATA4 expression.

**b.** Immunostaining for GFP (GATA4) and Krt7 in the esophagus after 2 weeks of Dox induction, following either 2 or 6 weeks of TAM induction in K5CreER:Rosa-rtTA:SmoM2:GATA4 (K5:Smo:R:GATA4) mice.

**c.** Experimental design showing 6 weeks of GATA4 induction following 6 weeks of Hedgehog pathway activation in K5:Smo:R:GATA4 mice.

**d.** Immunostaining for GFP (GATA4) and Krt14 in the esophagus in the condition described in (c).

**e.** Immunostaining for GFP (GATA4) and Cldn18 in the esophagus in the condition described in (c).

**f.** Immunostaining for GFP (GATA4) and Krt8 in the lesser curvature of the stomach in the condition described in (c).

**g.** Immunostaining for GFP (GATA4), p63, and Krt8 in the distal esophagus, 14 weeks after Dox treatment initiation following 1 week of TAM induction in K5:Smo:R:GATA4 mice.

SCJ, squamocolumnar junction; Fst, forestomach; Eso, esophagus; Stom, stomach; Col, columnar; Sq, squamous; Lum, lumen; LP, lamina propria; Hedgehog, HH; Dox, Doxycycline; TAM, tamoxifen.

Nuclei are counterstained with Hoechst (blue) in all images. Scale bar: 50 μm unless otherwise indicated.

### Extended data Fig.10

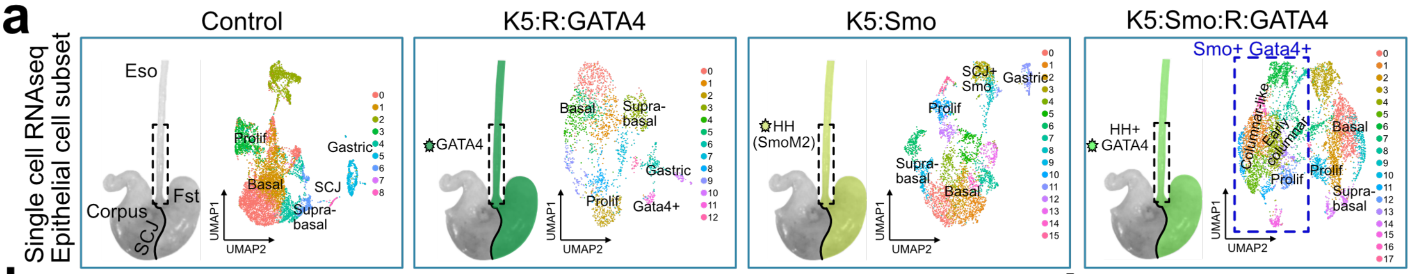

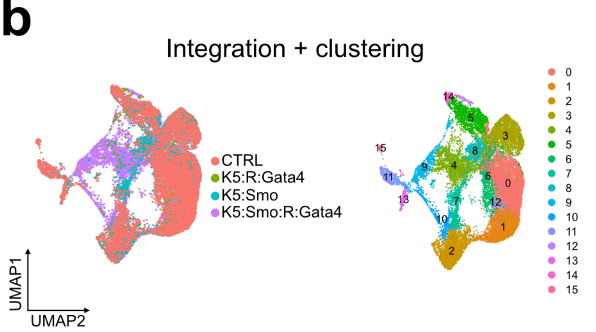

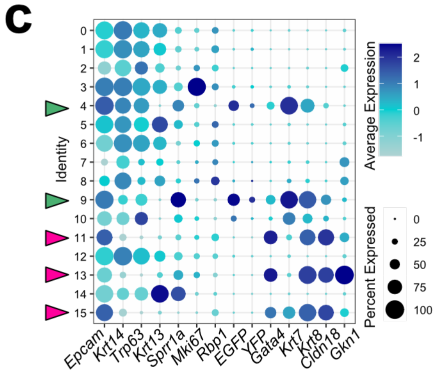

d

| | Nbr of cells in: | | | | |
|---|---|---|---|---|---|
| Ident | CTRL | K5:R:GATA4 | K5:Smo | K5:Smo:R:GATA4 | Tot |
| 0 | 2468 | 554 | 922 | 730 | 4674 |
| 1 | 1287 | 318 | 314 | 788 | 2707 |
| 2 | 817 | 262 | 814 | 662 | 2555 |
| 3 | 941 | 385 | 432 | 500 | 2258 |
| 4 | 24 | 39 | 43 | 1474 | 1580 |
| 5 | 687 | 221 | 303 | 324 | 1535 |
| 6 | 589 | 83 | 239 | 160 | 1071 |
| 7 | 226 | 20 | 530 | 232 | 1008 |
| 8 | 87 | 86 | 477 | 180 | 830 |
| 9 | 31 | 19 | 17 | 598 | 665 |
| 10 | 34 | 9 | 17 | 425 | 485 |
| 11 | 298 | 17 | 113 | 10 | 438 |
| 12 | 73 | 87 | 20 | 128 | 308 |
| 13 | 112 | 35 | 82 | 7 | 236 |
| 14 | 133 | 32 | 21 | 23 | 209 |
| 15 | 85 | 10 | 20 | 6 | 121 |

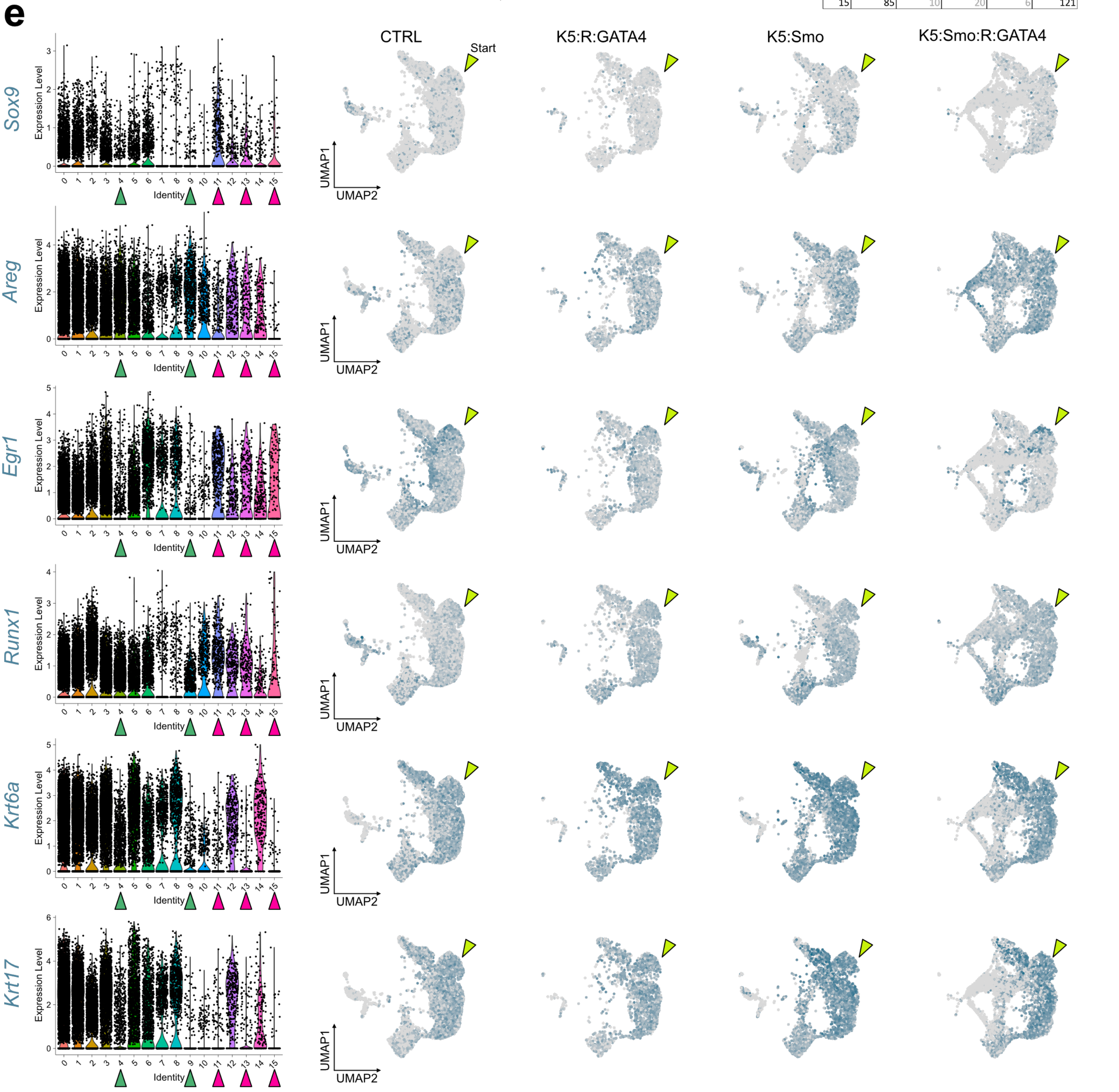

**Extended Data Fig. 10**
**a.** UMAP of the epithelial-cell subset extracted from the esophageal scRNA-seq dataset of CTRL, K5:R:GATA4, K5:Smo, and K5:Smo:R:GATA4 mice.
**b.** UMAP of the integrated dataset described in (a).
**c.** Dot plot depicting the expression of selected genes across the clusters in (b). Green arrows highlight Smo+ GATA4+ clusters, while fuchsia arrows highlight gastric clusters.
**d.** Table summarizing the number of cells per cluster and per condition. Values are dispalyed in grey when there are fewer than 30 cells.
**e.** Violin plot (left) and Feature plots (right) showing expression of the indicated mRNAs across the four conditions.

### Extended Data Fig. 11

**a**

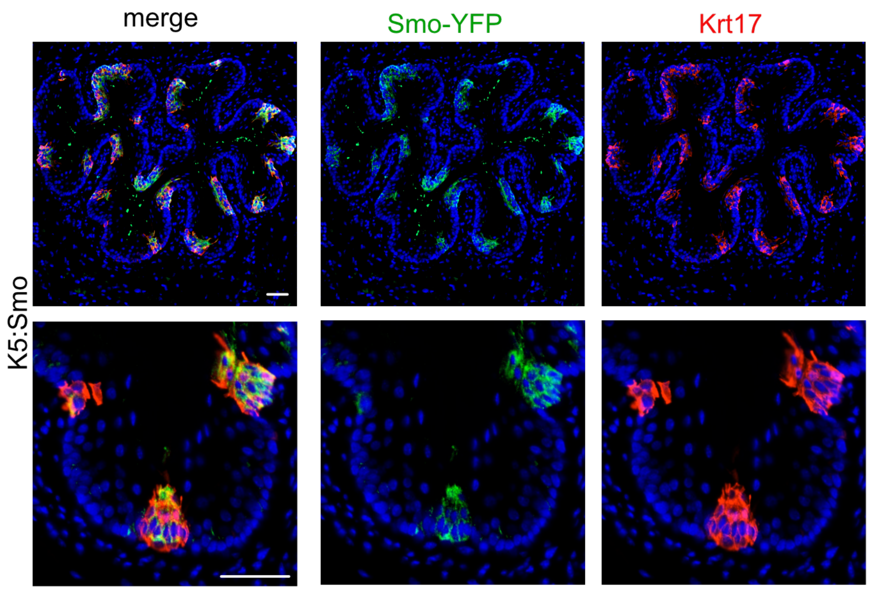

**b**

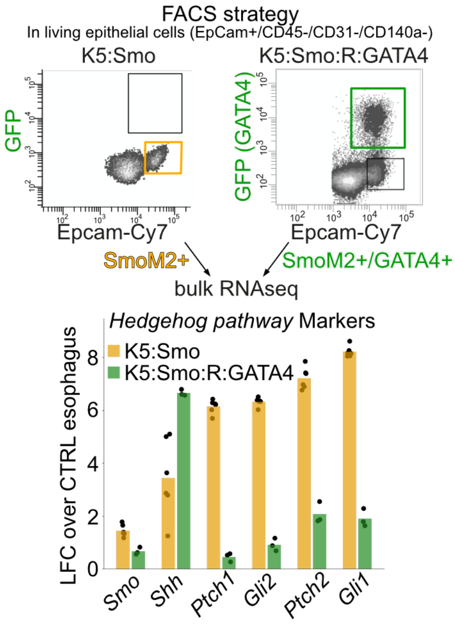

**c**

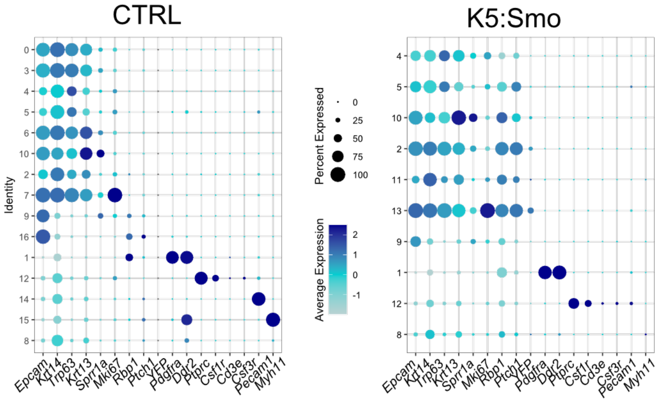

**d**

Number of cells per cluster

| ID | CTRL | K5:Smo | Tot | Annotation |
|---|---|---|---|---|
| 0 | 2536 | 12 | 2548 | Epith bas1 |
| 3 | 1539 | 1 | 1540 | Epith bas2 |
| 6 | 858 | 3 | 861 | Epith Suprabas2 |
| 7 | 859 | 1 | 860 | Epith prolif1 |
| 16 | 70 | 5 | 75 | Gastric2 |
| 14 | 281 | 21 | 302 | Endoth |
| 15 | 131 | 20 | 151 | Musc lis |
| 11 | 26 | 495 | 521 | Epith Cycling |
| 13 | 19 | 353 | 372 | Epith prolif Smo+ |
| 4 | 1006 | 41 | 1047 | Epith bas3 |
| 5 | 44 | 900 | 944 | Epith Basal Smo+ |
| 10 | 376 | 195 | 571 | Epith Suprabas1 |
| 2 | 66 | 1662 | 1728 | Epith Suprabas Smo+ |
| 9 | 485 | 255 | 740 | Gastric1 |
| 1 | 1768 | 472 | 2240 | Fibro |
| 12 | 301 | 84 | 385 | Immu |
| 8 | 244 | 531 | 775 | Dying cells |

**e**

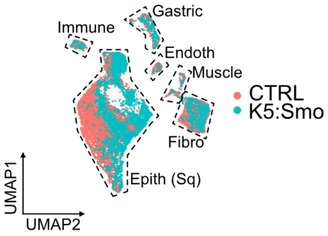

**Extended Data Fig. 11**

**a.** Immunofluorescence showing Smo-YFP and Krt17 expression in the esophagus of K5:Smo mouse. Nuclei are counterstained with Hoechst (blue) in all images. Scale bar: 50 µm.

**b.** FACS strategy : SmoM2+ cells (Epcam high) were isolated from K5:Smo mice 12 weeks after TAM-induced Hedgehog activation (as previously described[15]), and SmoM2+/GATA4+ cells (Epcam high GFP+) were isolated from K5:Smo:R:GATA4 mice in which GATA4 expression was induced by Doxycycline treatment for 6 weeks, initiated 6 weeks after TAM injection. Bulk RNA-seq on sorted cell populations was used to compare Hedgehog pathway gene expression. Log2 fold changes were calculated relative to CTRL esophageal epithelial cells.

**c.** Dot plot showing the expression of selected marker genes used for cluster annotation in the scRNA-seq dataset from CTRL and K5:Smo mice.

**d.** Table summarizing the number of cells per cluster and per condition, along with the corresponding cluster annotation. Values are shown in grey when fewer than 30 cells are present.

**e.** UMAP projection of cells from the merged CTRL and K5:Smo datasets, colored by experimental condition.

TAM, tamoxifen.

### Extended Data Fig. 12

**a**

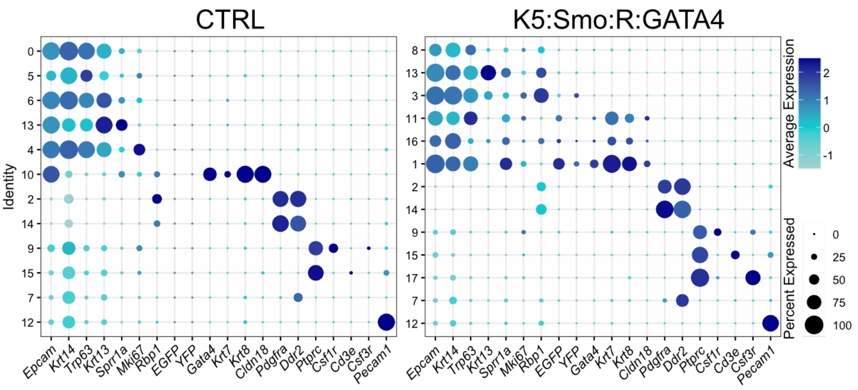

**b**

Number of cells per cluster

| ID | CTRL | K5:Smo:R:GATA4 | Annotation |
|---|---|---|---|
| 0 | 3284 | 0 | Epith bas1 |
| 5 | 1071 | 5 | Epith bas2 |
| 6 | 929 | 6 | Epith Suprab |
| 4 | 1827 | 0 | Epith bas prolif |
| 10 | 543 | 23 | Gastric |
| 8 | 3 | 762 | Epith bas Smo1 |
| 3 | 0 | 2029 | Epith bas prolif |
| 16 | 1 | 203 | Early metaplastic-like |
| 11 | 0 | 432 | Transitional squam |
| 1 | 5 | 2588 | Metaplastic-like |
| 17 | 1 | 156 | Neutrophils |
| 13 | 192 | 209 | Epith Suprab |
| 2 | 1657 | 640 | Fibro1 |
| 14 | 102 | 270 | Fibro2 |
| 9 | 265 | 441 | Macrophages |
| 15 | 77 | 232 | Lymphocytes |
| 7 | 368 | 451 | Muscu |
| 12 | 284 | 120 | Endoth |

**c**

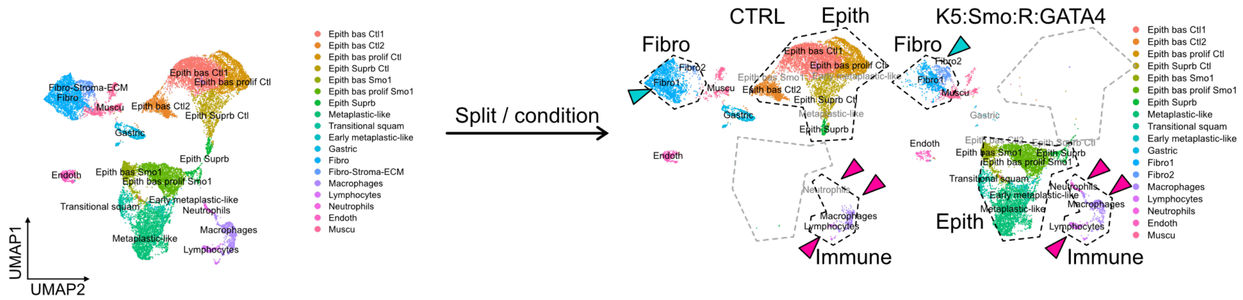

**d**

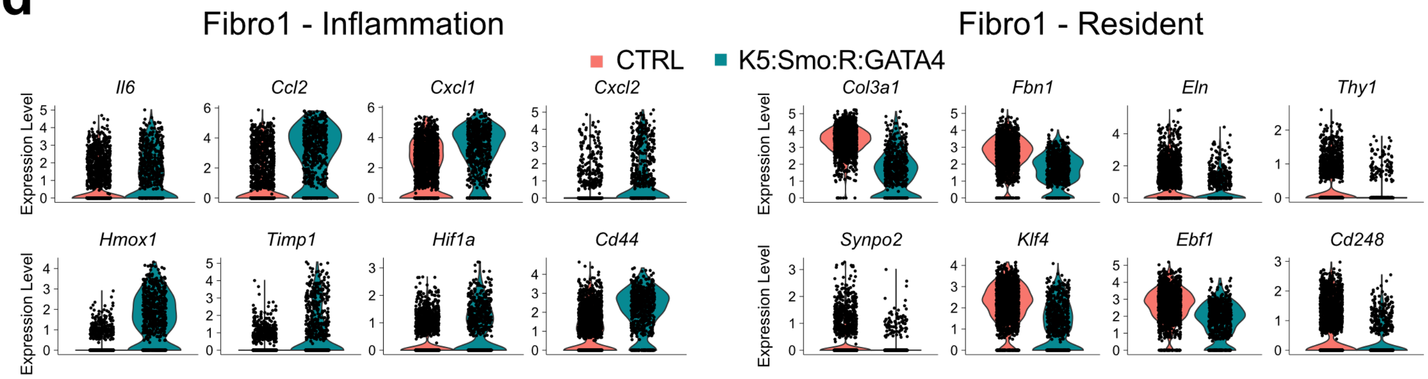

**e**

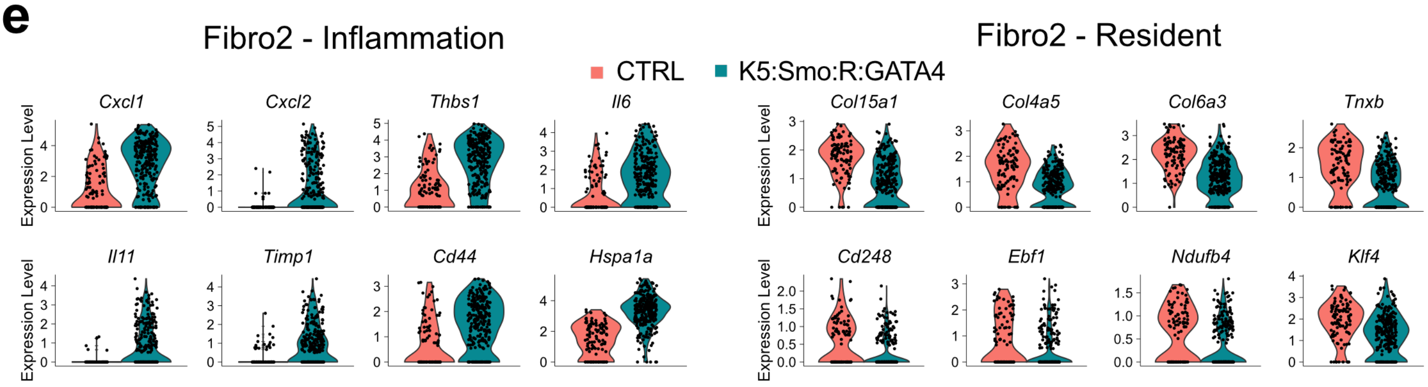

**Extended Data Fig. 12**

**a.** Dot plot showing the expression of selected marker genes used for cluster annotation in the scRNA-seq dataset from CTRL and K5:Smo:R:GATA4 mice.

**b.** Table summarizing the number of cells per cluster and per condition, along with the corresponding cluster annotation. Values are shown in grey when fewer than 30 cells are present.

**c.** Annotated UMAP of merged CTRL and K5:Smo:R:GATA4 datasets, shown as a combined view and split by condition. Blue arrows indicate fibroblast clusters; fuchsia arrows indicate immune clusters.

**d.** Violin plots showing the expression of inflammation- and quiescence-related markers in the Fibro1 cluster from CTRL and K5:Smo:R:GATA4 conditions.

**e.** Violin plots showing the expression of inflammation- and tissue-residency markers in the Fibro2 cluster from CTRL and K5:Smo:R:GATA4.

### Extended Data Fig. 13

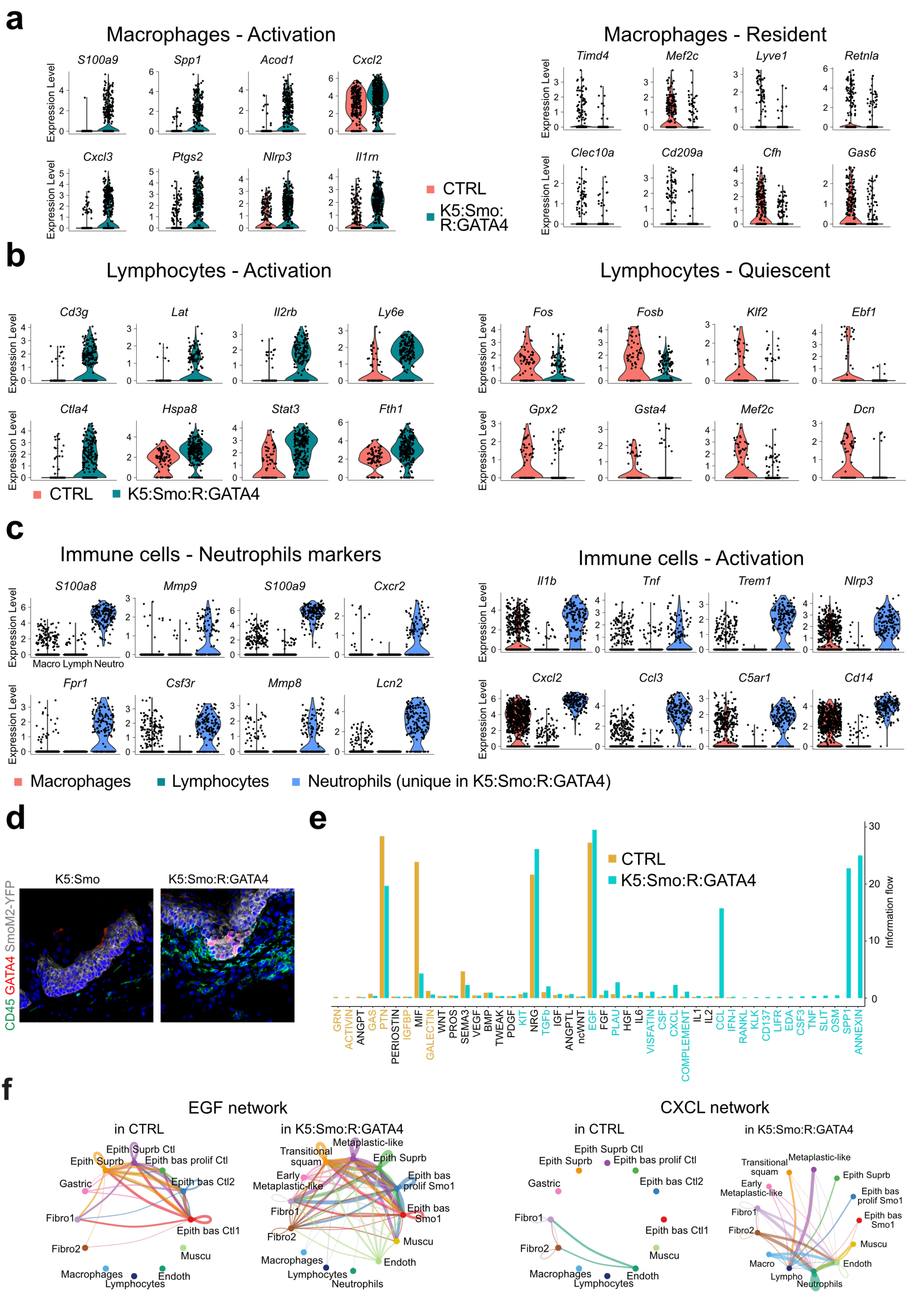

**Extended Data Fig. 13**

**a.** Violin plots showing the expression of activation- and tissue-residency markers in the macrophage cluster from CTRL and K5:Smo:R:GATA4 conditions.

**b.** Violin plots showing the expression of activation- and quiescence-related markers in the lymphocyte cluster from K5:Smo:R:GATA4 compared to CTRL.

**c.** Violin plots of neutrophil and activation marker expression across immune cell clusters from the merged CTRL and K5:Smo:R:GATA4 dataset.

**d.** Immunofluorescence staining for CD45, GATA4, and Smo-YFP in the esophagus of K5:Smo and K5:Smo:R:GATA4 mice.

**e.** Relative information flow quantified by *CellChat* in CTRL and K5:Smo:R:GATA4 esophageal cells, revealing alterations in intercellular signaling associated with co-expression of SmoM2 and GATA4 in the epithelium. Non-significant pathways are shown in black.

**f.** Circle plots illustrating changes in EGF and CXCL signaling networks in K5:Smo:R:GATA4 mouse esophagus compared to CTRL.

**Table S1**

Full linearized pTRE-β-globulin-Gata4-3×HA-IRES-GFP-polyA DNA sequence injected for transgenesis :

TTTACTCCCTATCAGTGATAGAGAACGTATGTCGAGTTTACTCCCTATCAGTGATAGAGAACGATGTCGAGTTTACT
CCCTATCAGTGATAGAGAACGTATGTCGAGTTTACTCCCTATCAGTGATAGAGAACGTATGTCGAGTTTACTCCCTA
TCAGTGATAGAGAACGTATGTCGAGTTTATCCCTATCAGTGATAGAGAACGTATGTCGAGTTTACTCCCTATCAGTG
ATAGAGAACGTATGTCGAGGTAGGCGTGTACGGTGGGAGGCCTATATAAGCAGAGCTCGTTTAGTGAACCGTCA
GATCGCCTGGAGAATTCTGCAGGTGAGTTTGGGGACCCTTGATTGTTCTTTCTTTTTCGCTATTGTAAAATTCATG
TTATATGGAGGGGGCAAAGTTTTCAGGGTGTTGTTTAGAATGGGAAGATGTCCCTTGTATCACCATGGACCCTCA
TGATAATTTTGTTTCTTTCACTTTCTACTCTGTTGACAACCATTGTCTCCTCTTATTTTCTTTTCATTTTCTGTAACTTT
TTCGTTAAACTTTAGCTTGCATTTGTAACGAATTTTTAAATTCACTTTTGTTTATTTGTCAGATTGTAAGTACTTTCTC
TAATCACTTTTTTTTCAAGGCAATCAGGGTATATTATATTGTACTTCAGCACAGTTTTAGAGAACAATTGTTATAATT
AAATGATAAGGTAGAATATTTCTGCATATAAATTCTGGCTGGCGTGGAAATATTCTTATTGGTAGAAACAACTACAT
CCTGGTCATCATCCTGCCTTTCTCTTTATGGTTACAATGATATACACTGTTTGAGATGAGGATAAAATACTCTGAGTC
CAAACCGGGCCCCTCTGCTAACCATGTTCATGCCTTCTTCTTTTTCCTACAGGCTAGCGCCGCCATGTACCAAAGC
CTGGCCATGGCCGCCAACCACGGCCCCCCGCCCGGCGCCTACGAAGCAGGTGGCCCTGGCGCCTTCATGCACA
GCGCGGGCGCCGCGTCCTCGCCCGTCTACGTGCCCACTCCACGAGTGCCGTCCTCTGTGCTGGGCCTGTCCTACC
TGCAGGGCGGTGGCAGTGCCGCTGCAGCTGGAACCACCTCGGGTGGCAGCTCCGGGGCCGGCCCGTCGGGTG
CAGGGCCTGGGACCCAGCAGGGTAGCCCTGGCTGGAGCCAAGCTGGAGCCGAGGGAGCCGCCTACACCCCGC
CGCCCGTGTCCCCGCGCTTCTCTTTCCCGGGGACTACTGGGTCCCTGGCGGCCGCTGCCGCCGCTGCCGCAGCC
CGGGAAGCTGCAGCCTACGGCAGTGGCGGCGGGGCGGCGGGCGCTGGTCTGGCTGGCCGAGAGCAGTACGG
GCGTCCGGGCTTCGCCGGCTCCTACTCCAGCCCCTACCCAGCCTACATGGCCGACGTGGGAGCATCCTGGGCCG
CAGCCGCTGCCGCCTCTGCCGGCCCCTTCGACAGCCCAGTCCTGCACAGCCTGCCTGGACGGGCCAACCCTGGA
AGACACCCCAATCTCGTAGATATGTTTGATGACTTCTCAGAAGGCAGAGAGTGTGTCAATTGTGGGGCCATGTCC
ACCCCACTCTGGAGGCGAGATGGGACGGACACTACCTGTGCAATGCCTGTGGCCTCTATCACAAGATGAACGG
CATCAACCGGCCCCTCATTAAGCCTCAGCGCCGCCTGTCCGCTTCCCGCCGGGTAGGCCTCTCCTGTGCCAACTG
CCAGACTACCACCACCACGCTGTGGCGTCGTAATGCCGAGGGTGAGCCTGTATGTAATGCCTGCGGCCTCTACAT
GAAGCTCCATGGGGTTCCCAGGCCTCTTGCAATGCGGAAGGAGGGGATTCAAACCAGAAAACGGAAGCCCAA
GAACCTGAATAAATCTAAGACGCCAGCAGGTCCTGCTGGTGAGACCCTCCCTCCCTCCAGTGGTGCCTCCAGCG
GTAACTCCAGCAATGCCACTAGCAGCAGCAGCAGCAGTGAAGAGATGCGCCCCATCAAGACAGAGCCCGGGCT
GTCATCTCACTATGGGCACAGCAGCTCCATGTCCCAGACATTCAGTACTGTGTCCGGCCACGGGCCCTCCATCCAT
CCAGTGCTGTCTGCTCTGAAGCTGTCCCCACAAGGCTATGCATCTCCTGTCACTCAGACATCGCAGGCCAGCTCC
AAGCAGGACTCTTGGAACAGCCTGGTCCTGGCTGACAGTCATGGGGACATAATCACCGCAGGAGGTGGAGGTG
GAGGTTACCCATACGATGTTCCAGATTACGCTGGATACCCATACGATGTTCCAGATTACGCTGGATACCCATACGAT
GTTCCAGATTACGCTTAATAGCCGCGGGCCCGGGATCCGCCCCTCTCCCTCCCCCCCCCCTAACGTTACTGGCCGA
AGCCGCTTGGAATAAGGCCGGTGTGCGTTTGTCTATATGTTATTTTCCACCATATTGCCGTCTTTTGGCAATGTGAG
GGCCCGGAAACCTGGCCCTGTCTTCTTGACGAGCATTCCTAGGGGTCTTTCCCCTCTCGCCAAAGGAATGCAAG
GTCTGTTGAATGTCGTGAAGGAAGCAGTTCCTCTGGAAGCTTCTTGAAGACAAACAACGTCTGTAGCGACCCTT
TGCAGGCAGCGGAACCCCCCACCTGGCGACAGGTGCCTCTGCGGCCAAAAGCCACGTGTATAAGATACACCTGC
AAAGGCGGCACAACCCCAGTGCCACGTTGTGAGTTGGATAGTTGTGGAAAGAGTCAAATGGCTCTCCTCAAGCG
TATTCAACAAGGGGCTGAAGGATGCCCAGAAGGTACCCCATTGTATGGGATCTGATCTGGGGCCTCGGTACACAT
GCTTTACATGTGTTTAGTCGAGGTTAAAAAAACGTCTAGGCCCCCCGAACCACGGGGACGTGGTTTTCCTTTGAA
AAACACGATGATAATATGGCCACAACCATGGTGAGCAAGGGCGAGGAGCTGTTCACCGGGGTGGTGCCCATCCT
GGTCGAGCTGGACGGCGACGTAAACGGCCACAAGTTCAGCGTGTCCGGCGAGGGCGAGGGCGATGCCACCTA
CGGCAAGCTGACCCTGAAGTTCATCTGCACCACCGGCAAGCTGCCCGTGCCCTGGCCCACCCTCGTGACCACCC
TGACCTACGGCGTGCAGTGCTTCAGCCGCTACCCCGACCACATGAAGCAGCACGACTTCTTCAAGTCCGCCATGC
CCGAAGGCTACGTCCAGGAGCGCACCATCTTCTTCAAGGACGACGGCAACTACAAGACCCGCGCCGAGGTGAA
GTTCGAGGGCGACACCCTGGTGAACCGCATCGAGCTGAAGGGCATCGACTTCAAGGAGGACGGCAACATCCTG

GGGCACAAGCTGGAGTACAACTACAACAGCCACAACGTCTATATCATGGCCGACAAGCAGAAGAACGGCATCAA
GGTGAACTTCAAGATCCGCCACAACATCGAGGACGGCAGCGTGCAGCTCGCCGACCACTACCAGCAGAACACC
CCCATCGGCGACGGCCCCGTGCTGCTGCCCGACAACCACTACCTGAGCACCCAGTCCGCCCTGAGCAAAGACCC
CAACGAGAAGCGCGATCACATGGTCCTGCTGGAGTTCGTGACCGCCGCCGGGATCACTCTCGGCATGGACGAGC
TGTACAAGTAAAGCGGCCGCATCGATAAGCTTGTCGACGATATCTCTAGAGGATCATAATCAGCCATACCACATTT
GTAGAGGTTTTACTTGCTTTAAAAAACCTCCCACACCTCCCCCTGAACCTGAAACATAAAATGAATGCAATTGTTG
TTGTTAACTTGTTTATTGCAGCTTATAATGGTTACAAATAAAGCAATAGCATCACAAATTTCACAAATAAAGCATTTT
TTTCACTGC

**Table S2**

Resource Table

| REAGENT or RESOURCE | SOURCE | IDENTIFIER |
|---|---|---|
| Antibodies | | |
| Chicken polyclonal anti-Krt14 1/10000 | BioLegend | 906004 |
| Rabbit monoclonal anti-Cldn18 1/2000 | abcam | ab203563 |
| Rat monoclonal anti-EpCam 1/500 | BioLegend | 118 202 ; RRID:AB_1089027 |
| Goat polyclonal anti-GFP 1/1000 | abcam | ab6673 ;RRID AB_305643 |
| Rabbit polyclonal anti-GFP 1/2000 | abcam | ab6556 ; RRID:AB_305564 |
| Rabbit monoclonal anti-Krt7 1/1000 | abcam | ab181598 |
| Rat monoclonal anti-Krt8 1/250 | DSHB | TROMA-I ; RRID:AB_531826 |
| Rabbit monoclonal anti-p63 1/1000 | abcam | ab124762 ; RRID:AB_10971840 |
| Rabbit Krt17 1/1000 | Cell Signaling technology | 4543 |
| Rabbit Cdx2 1/500 | abcam | ab76541 |
| Alexa Fluor 488 donkey anti-chicken | Jackson Immunoresearch | 703-545-155 ; RRID:AB_2340375 |
| Alexa Fluor 488 donkey anti-rabbit | Jackson Immunoresearch | 711-545-152 ; RRID:AB_2313584 |
| Alexa Fluor 488 donkey anti-goat | Jackson Immunoresearch | 705-545-147 ; RRID:AB_2336933 |
| Rhodamine Red X donkey anti-chicken | Jackson Immunoresearch | 703-295-155 ; RRID:AB_2340371 |
| Rhodamine Red X donkey anti-rabbit | Jackson Immunoresearch | 711-295-152 ; RRID:AB_2340613 711-297-003 ; RRID:AB_2340615 |
| Alexa Fluor 647 donkey anti-chicken | Jackson Immunoresearch | 703-605-155 ; RRID:AB_2340379 |
| Alexa Fluor 647 donkey anti-rat | Jackson Immunoresearch | 712-605-153 ; RRID:AB_2340694 |
| Rat anti-CD45 PE | BioLegend | 103106 ; RRID:AB_312971 |
| Rat anti-CD140a PE | BioLegend | 135906 ; RRID:AB_1953269 |
| Rat anti-CD31 PE | BioLegend | 102508 ; RRID:AB_312915 |
| Rat anti-CD326(EpCam) APC-Cy7 | BioLegend | 118218 ; RRID:AB_2098648 |
| Chemicals, Peptides, and Recombinant Proteins | | |
| Bovin Serum Albumin | Capricorn scientific | BSA-IT Lot CP14-1028 |
| Triton X-100 | Sigma-Aldrich | T8787 |
| Glycergel | Dako | C0563 |
| 1,4-Diazabicyclo[2,2,8]octane (Dabco) | Sigma-Aldrich | D27802 |
| Hoechst 33258 (10mg/mL solution) | Molecular probes | Cat # H3569 |
| Tamoxifen | A&E Scientific | 10540-29-1 |
| Sunflower | Sigma-Aldrich | S5007 |
| Formaldehyde 4% | VWR | 116 994 55 |
| Collagenase I | A.G. Scientific | C-2823 |

| | | |
|---|---|---|
| O.C.T. | Tissue-Tek | Cat#4583 |
| Horse Serum | Capricorn scientific | HOS-1b |
| Foetal Bovine Serum | Gibco | Cat# 10270-106 |
| Trypsin solution 2.5% | A&E Scientific | TRY-2B10 |
| Sucrose/ Saccharose 30% in PBS | Milipore Merck | 1076511000 |
| Critical Commercial Assays | | |
| E.Z.N.A Total RNA Kit | Omega BIO-TEK | SKU: R6834-01 |
| 2100 Bioanalyzer | Agilent | G293BA) |
| MinElute purification kit | QIAGEN | Cat # 28004 |
| Ovation SoLo RNA-Seq System | NuGEN | Part # 0501-32 |
| NEBNext High-Fidelity 2x PCR Master Mix | New England BioLabs | M0541S |
| Experimental Models: Organisms/Strains | | |
| K5-CreER$^{T2}$ knock-in<br>B6N.129S6(Cg)-Krt5<tm1.1(cre/ERT2) Blh>/J | The Jackson Laboratory | Stock# 029155 ;<br>RRID:IMSR_JAX:029155 |
| R26SmoM2<br>Gt(ROSA)26Sor<tm1(Smo/EYFP) Amc>/J | The Jackson Laboratory | Stock# 005130 ;<br>RRID:IMSR_JAX:005130 |
| TetO-GATA4-IRES-GFP | Benjamin Beck Laboratory | N/A |
| Rosa26-CAGs-LSL-rtTA3 | The Jackson Laboratory | Stock#029617, RRID: IMSR_JAX:029617 |
| Software and Algorithms | | |
| Zen Blue | Zeiss | N/A |
| FACSDiva | BD Biosciences | N/A |
| STAR version 2.5.3a | Alexander Dobin (2009-2019) | https://github.com/alexdobin/STAR<br>PMID:23104886<br>DOI:10.1093/bioinformatics/bts635 |
| Degust 4.1.183: *interactive RNA-seq analysis* | David R. Powell | http://degust.erc.monash.edu/ |
| R version 4.4.2 or 4.3.1 | GNU project | https://cran.r-project.org/ |
| Bioconductor | | https://www.bioconductor.org/install/ |
| Cell Ranger (version 3.1.0) | 10x Genomics | https://www.10xgenomics.com/support/software/cell-ranger/latest |
| Other | | |
| Mouse reference genome GRCm38/mm10 | Genome Reference Consortium | https://www.ncbi.nlm.nih.gov/grc/mouse |
| C5 collection MSigDB for mouse | Walter+Eliza Hall Bioinformatic Resources | http://bioinf.wehi.edu.au/software/MSigDB/ |
| Human mircroarray data on Barrett samples and normal squamous samples #1 | GEO : Hyland et al. | GSE39491 |
| Human mircroarray data on Barrett samples and normal squamous samples #2 | GEO : di Pietro et al. | GSE34619 |
| Human mircroarray data on Barrett samples and normal squamous samples #3 | GEO : Ostrowski et al. | GSE36223 |

**Table S3**

**Single Cell RNAseq parameters**

| Object used in the main figures | | | | | | | | | |
|---|---|---|---|---|---|---|---|---|---|
| Figure | Samples | Treatment | Combination | Nbr of cells before subset | Subset | Nbr of cells after subset | npcs | resolution | alias |
| 3a | Lesser and Greater curvatures from CTRL and K5:Smo:R:GATA4 stomach | DOX 14 weeks following TAM 1 week | merge of objects CtlPC.Rds, SmGPC.Rds, CtlGC.Rds, SmGGC.Rds | 20885 | / | 20885 | 19 | 0.6 | CtlsSmGsMrg.Rds |
| 3b-h | Lesser curvature from CTRL and K5:Smo:R:GATA4 stomach | DOX 14 weeks following TAM 1 week | merge of objects CtlPC.Rds, SmGPC.Rds | 13011 | / | 13011 | 9 | 0.6 | PCs-res0.6.Rds |
| 5a-f | Esophagus K5:Smo:R:GATA4 | DOX 6 weeks following TAM 6 weeks | single object SmoGata4.Rds | 8567 | remove clusters: 2,6,11,12,4,13 | 6243 | 12 | 1.2 | SmGata4Sub1.Rds |
| 5g-h | Esophagus CTRL, K5:R:GATA4, K5:Smo, K5:Smo:R:GATA4 subset epithelial cells | DOX 6 weeks following TAM 6 weeks | integration RPCA of objects CtlG.Rds, Gata4.Rds, SmoG.Rds, SmoGata4.Rds to give rise to Gata4AllInteg3.Rds | 26843 | remove clusters: 1,15,16,14,13,7, 18 | 20680 | 16 | 0.45 | Integ3rpcaSub.Rds |
| 5i | Esophagus CTRL, K5:Smo | DOX 6 weeks following TAM 6 weeks | merge of objects CtlG.Rds, SmoG.Rds | 15660 | / | 15660 | 13 | 0.45 | CtlSmoMerg.rds |

| Object used in the Extended figures | | | | | | | | | |
|---|---|---|---|---|---|---|---|---|---|
| Figure | Samples | Treatment | combination | Nbr of cells before subset | Subset | Nbr of cells after subset | npcs | resolution | alias |
| S6 a-c | Lesser and Greater curvatures from CTRL and K5:Smo:R:GATA4 stomach | DOX 14 weeks following TAM 1 week | merge of objects CtlPC.Rds, SmGPC.Rds, CtlGC.Rds, SmGGC.Rds | 20885 | / | 20885 | 19 | 0.6 | CtlsSmGsMrg.Rds |
| S7a-f | Lesser curvature from CTRL and K5:Smo:R:GATA4 stomach | DOX 14 weeks following TAM 1 week | merge of objects CtlPC.Rds, SmGPC.Rds | 13011 | / | 13011 | 9 | 0.6 | PCs-res0.6.Rds |
| S10b | Esophagus CTRL, K5:R:GATA4, K5:Smo, K5:Smo:R:GATA4 subset epithelial cells | DOX 6 weeks following TAM 6 weeks | integration RPCA of objects CtlG.Rds, Gata4.Rds, SmoG.Rds, SmoGata4.Rds to give rise to Gata4AllInteg3.Rds | 26843 | remove clusters: 1,15,16,14,13,7, 18 | 20680 | 16 | 0.45 | Integ3rpcaSub.Rds |
| S11c-d | Esophagus CTRL, K5:Smo | DOX 6 weeks following TAM 6 weeks | merge of objects CtlG.Rds, SmoG.Rds | 15660 | / | 15660 | 13 | 0.45 | CtlSmoMerg.rds |
| S12-S13 | Esophagus CTRL + K5:Smo:R:GATA4 | DOX 6 weeks following TAM 6 weeks | merge of objects CtlG.Rds, SmoGata4.Rds | 19176 | / | 19176 | 00:00 | 0.45 | Ctl.SmoGata4Merg.rds |

Processed objects used for further analysis in the figures

| Samples | Treatment | combination | Nbr of cells before subset | Subset | Nbr of cells after subset | npcs | resolution | alias |
|---|---|---|---|---|---|---|---|---|
| Lesser curvature from CTRL stomach | DOX 14 weeks following TAM 1 week | single object | 6993 | 1000 < nCount_RNA <40000; percent.mt<20 | 6263 | 13 | 0.45 | CtlPC.Rds |
| Lesser curvature from K5:Smo:R:GATA4 stomach | DOX 14 weeks following TAM 1 week | single object | 8369 | 1000 < nCount_RNA <80000; percent.mt<60 | 6748 | 17 | 1 | SmGPC.Rds |
| Greater curvature from CTRL stomach | DOX 14 weeks following TAM 1 week | single object | 3504 | 1000 < nCount_RNA <40000; percent.mt<20 | 3128 | 20 | 0.45 | CtlGC.Rds |
| Greater curvature from K5:Smo:R:GATA4 stomach | DOX 14 weeks following TAM 1 week | single object | 5800 | 1000 < nCount_RNA <80000; percent.mt<40 | 4746 | 17 | 0.6 | SmGGC.Rds |
| Esophagus CTRL | DOX 6 weeks following TAM 6 weeks | single object | 13655 | 1000 < nCount_RNA <40000; percent.mt<40 | 10609 | 00:00 | 0.3 | CtlG.Rds |
| Esophagus K5:R:GATA4 | DOX 6 weeks following TAM 6 weeks | single object | 3044 | 1500 < nCount_RNA <100000; percent.mt<20 | 2616 | 00:00 | 0.3 | Gata4.Rds |

| | | | | | | | | |
|---|---|---|---|---|---|---|---|---|
| Esophagus K5:Smo | DOX 6 weeks following TAM 6 weeks | single object | 6256 | 700 < nCount_RNA <50000; percent.mt<50 | 5051 | 16 | 0.6 | SmoG.Rds |
| Esophagus K5:Smo:R:GATA4 | DOX 6 weeks following TAM 6 weeks | single object | 10477 | 800 < nCount_RNA <60000; percent.mt<40 | 8567 | 12 | 0.45 | SmoGata4.Rds |
| Esophagi CTRL, K5:R:GATA4, K5:Smo, K5:Smo:R:GATA4 | DOX 6 weeks following TAM 6 weeks | integration RPCA | 26843 | / | 26843 | 16 | 0.45 | Gata4AllInteg3.Rds |
